# Immune Cell Dysfunction of SARS-CoV-2: Mathematical Modeling of the Within-Host Immune Dynamics

**DOI:** 10.1101/2025.10.03.680253

**Authors:** Pagnapech Ngoun, Nicolas Alvarez, Ayesh Awad, Hwayeon Ryu

**Author notes:** Corresponding author (Hwayeon Ryu). Email addresses:* (Pagnapech Ngoun), (Nicolas Alvarez), (Ayesh Awad).

## Abstract

The COVID-19 pandemic has spurred extensive research into viral transmission and control, yet the mechanisms of the human immune response to SARS-CoV-2 remain incompletely understood, particularly the role of natural killer (NK) cells and cytokine regulation in disease severity. Mathematical modeling provides a powerful approach to bridge this gap by linking viral dynamics with immune interactions. In this work, we develop a mechanistic within-host model, formulated in a system of coupled ordinary and delayed differential equations, to investigate the contributions of NK cell activity, interferon signaling, and pro-inflammatory cytokines to viral clearance and disease outcome. Model parameters are estimated from experimental data, and computational simulations are used to explore how dysregulated NK responses and cytokine feedback loops may drive divergent clinical outcomes. Local sensitivity analysis identifies the most influential parameters shaping host–pathogen dynamics, highlighting potential control points for intervention. In addition, knockdown simulations are performed to mimic potential therapeutic interventions, allowing us to evaluate their advantages and limitations *in silico*. These findings provide mechanistic insights into COVID-19 immune dynamics and offer a foundation for guiding the design of future treatment strategies.

**Highlights:**

- We develop a within-host mathematical model of the SARS-CoV-2 immune response.
- The model incorporates key cytokines and immune cells including Natural Killer cells.
- Numerical simulations reproduce cytokine storms and NK cell dysfunction in severe disease.
- Sensitivity analysis identifies parameter impact and potential therapeutic interventions.

## 1. Introduction

Despite an extensive volume of research, the mechanisms governing host immune dynamics in COVID-19 remain incompletely understood. Severe disease is often associated with hyperinflammatory responses, yet many studies have relied on heterogeneous cohorts with limited longitudinal sampling and rarely integrate viral load, cytokine profiles, and immune-cell phenotypes within the same individuals across disease severity. Confounding factors such as vaccination status, prior infection, viral variants, and therapeutic interventions further complicate inference [1], while blood-based measurements may fail to capture immune activity in tissue compartments central to pathology [2, 3]. As a result, many findings remain correlative, leaving unresolved how cytokine programs shape immune cell transitions, which feedback loops govern disease resolution versus escalation, and when interventions are most effective [4, 5].

A hallmark pathology of severe COVID-19 is the cytokine storm, characterized by excessive cytokine release, immune cell over-activation, and tissue injury [6, 7]. Dysregulated innate and adaptive responses, including lymphopenia and overactivation of myeloid populations, contribute to lung injury, multi-organ failure, and poor prognosis [8, 9]. Among immune subsets, natural killer (NK) cell dysfunction has emerged as an important feature. Although NK cell numbers are generally preserved, their cytotoxic activity is impaired, often in the context of elevated interleukin-6 (IL-6) [10]. The underlying biology of this dysfunction remains poorly understood, and existing therapies, including corticosteroids [10], IL-6 inhibitors such as tocilizumab [11], and antivirals like Paxlovid [12], provide only partial benefit. These findings highlight the need for deeper mechanistic insight into NK biology and its integration within the broader immune response.

Mathematical modeling has provided valuable tools to explore host–virus interactions in COVID-19. Differential equation frameworks have been used to capture viral kinetics together with elements of the innate and adaptive immune response. Such models have reproduced observed clinical trajectories, including viral load dynamics, lymphocyte depletion, and cytokine production, while also generating mechanistic hypotheses to explain differences between mild and severe cases [13, 14, 15]. By explicitly linking viral replication with immune-mediated control, these efforts have underscored how the timing and coordination of immune responses strongly influence disease outcomes.

Among a series of more recent within-host modeling studies [16, 17, 18, 19, 20, 21], the ODE-based work for coupling viral replication with inflammatory responses [16, 22] have focused on therapeutic interventions. Their work showed that antiviral effectiveness depends critically on both potency and initiation time, and predicting scenarios of viral rebound under short-course early treatment. More studies [23, 24] incorporated dense, daily viral load sampling into mechanistic models to quantify heterogeneity in viral expansion, clearance, and peak timing, while also linking viral load trajectories to infectiousness. A more recent study [25] extended and unified these approaches by analyzing a large data set of viral loads, identifying six distinct shedding patterns and modeling how immune timing and intensity explain divergent peak loads, durations, and rebound phenomena. Collectively, these studies highlight how individual-level variation in immune and viral dynamics shapes both therapeutic outcomes and transmission potential, while emphasizing the central role of immune heterogeneity and treatment timing in controlling infection.

Despite these advances, natural killer (NK) cells are often absent from within-host models or represented only in highly simplified terms. One recent effort explicitly included NK cells [26], but considered only simplified cytokine-mediated dynamics, leaving NK cell dysfunction unexplored. Other models treat NK cells as generic innate effectors that aid early viral clearance, without accounting for their context-dependent impairment [15, 27]. Yet clinical studies suggest that NK cells, while numerically preserved in many COVID-19 patients, exhibit impaired cytotoxic function, particularly in the presence of elevated interleukin-6 (IL-6). The failure to capture these altered effector functions limits our ability to use models to probe how NK dysfunction contributes to uncontrolled inflammation or prolonged viral persistence. Likewise, the within-host studies (e.g., [16, 22, 23, 24, 25]) did not explicitly distinguish disease severity by immune or cytokine responses, largely due to limited experimental data. Incorporating NK dysfunction and cytokine-mediated pathways into a model could therefore provide new insights into the cellular feedback loops that drive divergent clinical trajectories between mild and severe disease.

In addition, cytokine signaling pathways remain incompletely represented in previous modeling frameworks. While type I interferons (IFN-*α* or IFN-*β*) are often included [14], type II IFN (IFN-*γ*) and TNF-*α* are rarely modeled despite evidence linking their dysregulation to severe outcomes [7, 28, 29, 30, 31, 32].

TNF-*α* is a pro-inflammatory mediator that recruits immune cells but can also exacerbate pathology [33, 34, 35, 36]. Elevated TNF-*α* levels are observed in severe COVID-19 [6, 7, 37], where it drives infiltration of NK cells, macrophages, neutrophils, and dendritic cells [35, 38]. Anti-TNF-*α* therapies have been proposed [38, 39] and shown to reduce hospitalization risk among infected patients [40], though their prophylactic use in uninfected populations remains uncertain and ethically challenging.

IFN-*γ* is another cytokine implicated in COVID-19 pathogenesis, with both antiviral and immunoregulatory functions [41, 42]. Although protective in many infectious diseases [43, 44] and retaining antiviral activity in COVID-19 [45], reports of IFN-*γ* levels are inconsistent across severities [7, 28, 29, 30]. Some studies link high IFN-*γ* to severe disease, others to diminished expression in critical cases, underscoring its complex role. In addition, a study [46] suggested that IFN-*γ* and TNF-*α* may act synergistically. They showed both trigger macrophage apoptosis, suggesting targeted inhibition could mitigate immunopathology. This synergy highlights a critical but underexplored mechanism of COVID-19 and underscores the need for models that jointly incorporate IFN-*γ* and TNF-*α*.

Here, we develop a mathematical model of SARS-CoV-2 infection that explicitly incorporates NK cytotoxic function and the interactions of TNF-*α* and IFN-*γ* with both innate and adaptive immune responses. Our goal is to move beyond correlative findings and generate mechanistic insights into how specific cytokine programs and immune subsets drive divergent clinical trajectories. By anchoring our framework to prior modeling efforts and clinical observations, we aim to clarify how NK cells and cytokine interactions contribute to immune dysregulation, to identify control points that tip the balance between viral clearance and immunopathology, and to provide testable predictions for the targeting of immunomodulatory therapies. Ultimately, this work seeks to advance mechanistic understanding of COVID-19 pathogenesis and inform future therapeutic strategies.

Our paper is structured as follows: Section 2 provides the description of mathematical model we developed (Sec. 2.1), the model equations (Sec. 2.2), and the parameter estimation and numerical simulations (Sec. 2.3). Section 3 provides our main results for mild versus severe disease (Sec. 3.1), for local sensitivity analysis (Sec. 3.2), for *in silico* knockdown analysis on selected immune cells from mild disease (Sec. 3.3), and on pro-inflammatory cytokines from severe case (Sec. 3.4). We discuss our main findings in a broader context, model limitations, and suggestions for future work in Section 4. The Supplementary Materials provide more details on the process of parameter estimation and homeostasis calculation, and additional model results.

## 2. Mathematical Model and Methods

### 2.1. Model description

To study the dynamics of SARS-CoV-2 infection and the immune cell dysfunctions responsible for severe disease case, we develop a mathematical model of host-pathogen interactions, adapted and substantially expanded from [14]. While the previous work of [14] focused on type I interferon (type I IFN) dynamics against COVID-19 virus and its relationship to disease severity, we additionally incorporate IFN-*γ*, TNF-*α*, and Natural Killer (NK) cells into our model due to their critical roles in immune response to COVID-19. IFN-*γ* is a pleiotropic cytokine, which acts as a potent antiviral agent, but can induce excessive macrophage apoptosis [47, 48]. TNF-*α* is a pro-inflammatory cytokine responsible for the up-regulated production of other pro-inflammatory cytokines [39] and the recruitment of NK cells. NK cell dysfunction is observed in severe cases of COVID-19, causing substantially less infected cells to be eliminated [49].

Figure 1 shows a schematic diagram of our model which consists of three main sets of components: lung epithelial cells, immune cells (both innate and adaptive), and cytokines. Susceptible lung epithelial cells (*S*) that encounter SARS-CoV-2 (*V*) may become infected (*I*), as shown in Fig. 1A. Upon infection, the immune response is orchestrated by cytokines that stimulate resident immune cell subsets and recruit additional cells from the bone marrow and circulation. Infected cells either die (*D*) due to viral cytopathic effects or immune-mediated killing by inflammatory macrophages (*M*_Φ*I*_), neutrophils (*N*), CD8^+^ T cells (*T*), or NK cells (*K*), or they produce new viruses through replication. Infected cells also secrete type I IFN [50], which induce resistance in lung epithelial cells (*R*) and thereby limit further viral infection [51].

**Figure 1.**
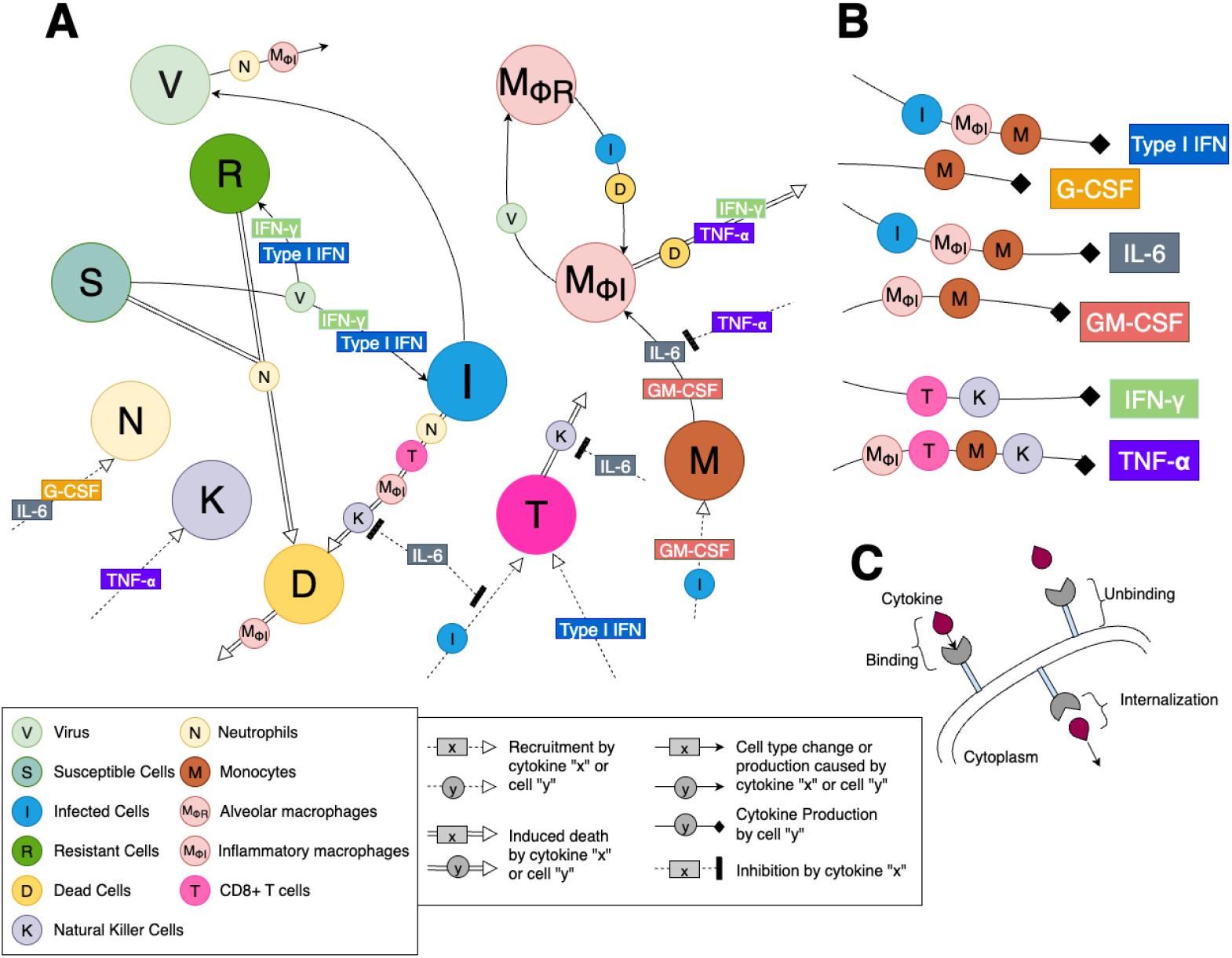
Schematic diagram of immune response to SARS-CoV-2 model. The model in Eqs. (1)–(23) is illustrated by the detailed dynamics for cells, cytokines, and their interactions, **B)** for cytokine production, and **C)** for cell-cytokine binding kinetics. Different lines represent cell recruitment (dotted line with triangle arrow), induced cell death (double line), cell type change or production (solid line with triangle arrow), cytokine production (solid line with rhombus arrow), and cytokine inhibition (dotted line with rectangle arrow). **A)** Virus (*V*) infects susceptible lung epithelial cells which can then turn into infected (*I*) or resistant (*R*) cells depending on the concentration of type I IFN and of IFN-*γ*. Infected cells can die and produce new virus (i.e., viral replication) or can be removed by inflammatory macrophages (*M*_Ф*I*_), CD8^+^ T cells (*T*), neutrophils (*N*) or NK cells (*K*) via apoptosis induction to become dead cells (*D*). NK cells eliminate target cells and are recruited by TNF-*α* while IL-6 can inhibit the cytotoxic activity of NK cells. Neutrophils can cause collateral damage (death) in all epithelial cells, including healthy ones, and are recruited by G-CSF and IL-6 concentrations. CD8^+^ T cells are recruited by infected cells and up-regulated by type I IFN concentration while the T cell recruitment being inhibited by IL-6 concentration. Monocytes (*M*) are recruited by infected cells and GM-CSF, then differentiate into inflammatory macrophages based on the concentrations of GM-CSF and IL-6. This differentiation process can be inhibited by TNF-*α*. Tissue-resident macrophages (*M*_F*R*_) also become inflammatory macrophages in response to the presence of dead and infected cells. The death of inflammatory macrophages can be induced by dead cells, TNF-*α*, and IFN-*γ*. Inflammatory macrophages are responsible for clearing up dead cells. Type I IFN is produced by infected cells, inflammatory macrophages, and monocytes while IFN-*γ* by T cells and NK cells. Monocytes are the sole producer of G-CSF. Both monocytes and inflammatory macrophages serve as producers for IL-6 and GM-CSF though IL-6 is also produced by infected cells. Inflammatory macrophages, T cells, monocytes, and NK cells all produce TNF-*α*. **C)** Each cell-cytokine interaction modeled includes cytokine receptor binding, internalization process, and unbinding kinetics.

The dynamics of inflammatory macrophages are governed by three distinct pathways: (i) conversion of tissueresident alveolar macrophages (*M*_Φ*R*_) in response to dead or infected cells [52], (ii) activation of alveolar macrophages by IFN-*γ* or TNF-*α* [53], and (iii) differentiation of monocytes (*M*) stimulated by IL-6, TNF-*α*, and GM-CSF [54]. Monocytes, recruited by infected cells and GM-CSF, therefore represent a key precursor population. Once differentiated, inflammatory macrophages clear dead cells but eventually undergo exhaustion (leading to cell death) during this process. Neutrophils (*N*) are recruited by IL-6 and G-CSF [55], and can induce collateral damage in both infected and uninfected epithelial cells [56, 57]. Their levels increase in COVID-19, further elevated by G-CSF and IL-6, which are strongly up-regulated in severe cases [6]. As part of the adaptive immune response, CD8^+^ T cells (*T*) are recruited to infection sites following antigen recognition, with their expansion modulated positively by type I IFN and negatively by IL-6 [58, 59, 60].

The production dynamics of cytokines are summarized in Fig. 1B. IFN-*γ* is primarily produced by T cells and NK cells, enacting antiviral mechanisms [47]. TNF-*α*, largely secreted by inflammatory macrophages and monocytes [61, 62], contributes to the excessive inflammation observed in severe patients [38]. Monocytes serve as major cytokine producers, generating all cytokines in the model except IFN-*γ* [63]. IL-6 not only drives monocyte-to-macrophage differentiation but also inhibits key processes, such as T cell recruitment by infected cells and NK cell–mediated clearance of infected cells [64]. G-CSF, produced solely by monocytes, further amplifies neutrophil responses. Cytokines function via receptor-mediated binding. Only free, unbound cytokines can signal effectively [65]; hence, our model explicitly distinguishes between bound and unbound cytokines, with receptor binding, internalization, and unbinding kinetics, as illustrated in Fig. 1C.

### 2.2. Model equations

Our model is formulated as a system of twenty-three coupled nonlinear ordinary and delay differential equations (Eqs. (1)–(23)) for eleven cell populations and six cytokines, in which, for each cytokine, both unbound and bound concentrations are modeled explicitly. The dynamics for lung epithelial cells are adapted from [14], except that IFN-*γ* now affects the infection and viral replication processes alongside type I IFN (see Fig. 1A). In addition, constants for each cytokine’s average receptor number used in the corresponding equations (i.e., Eqs. (12)–(23)) are provided in (A.1)–(A.6). Tables 1 and 2 list the variables and selected biological pathways represented in the equations.

**Table 1:**
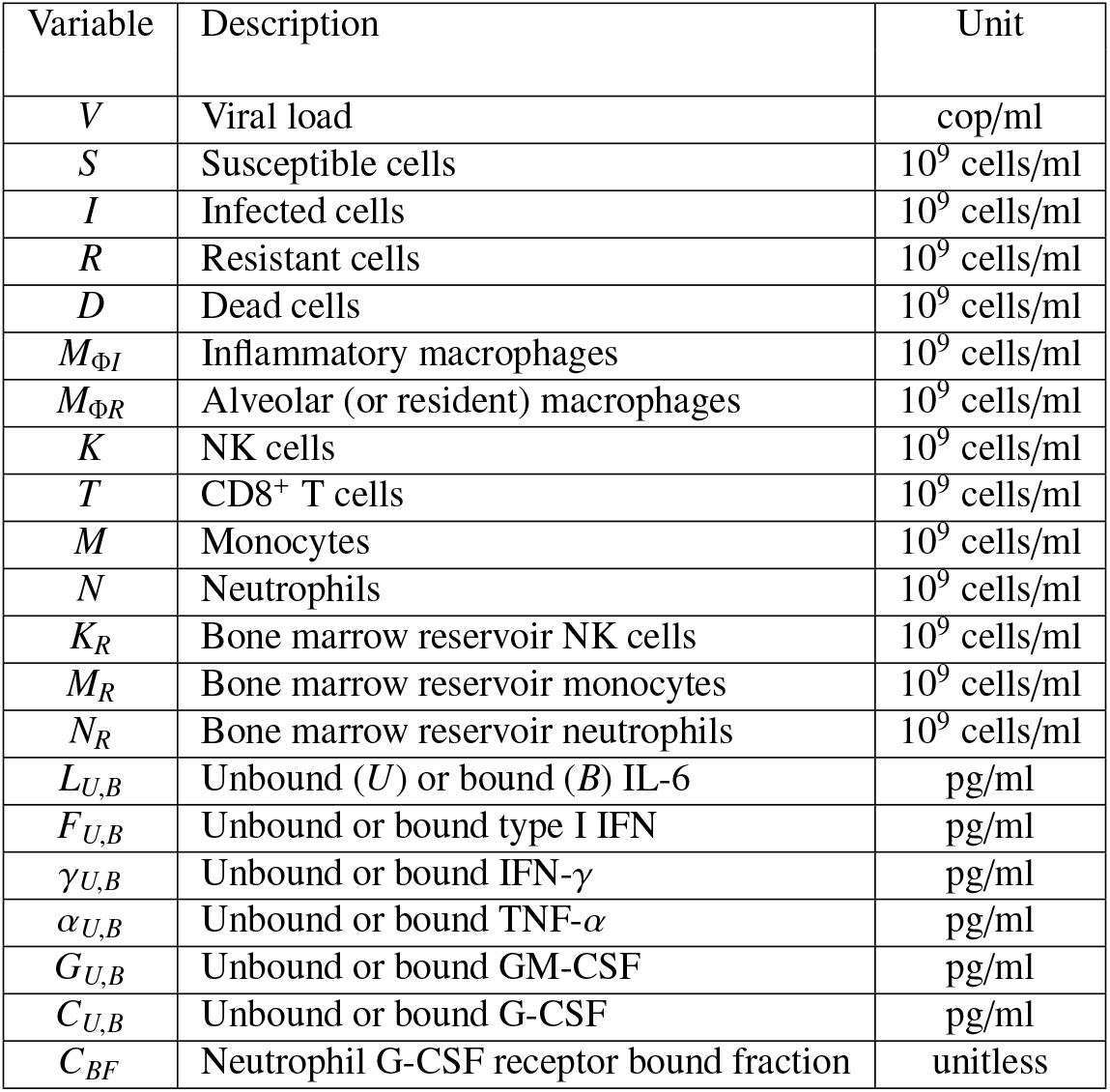
List of model variables and their description in Eqs. (1)–(23).

**Table 2:**
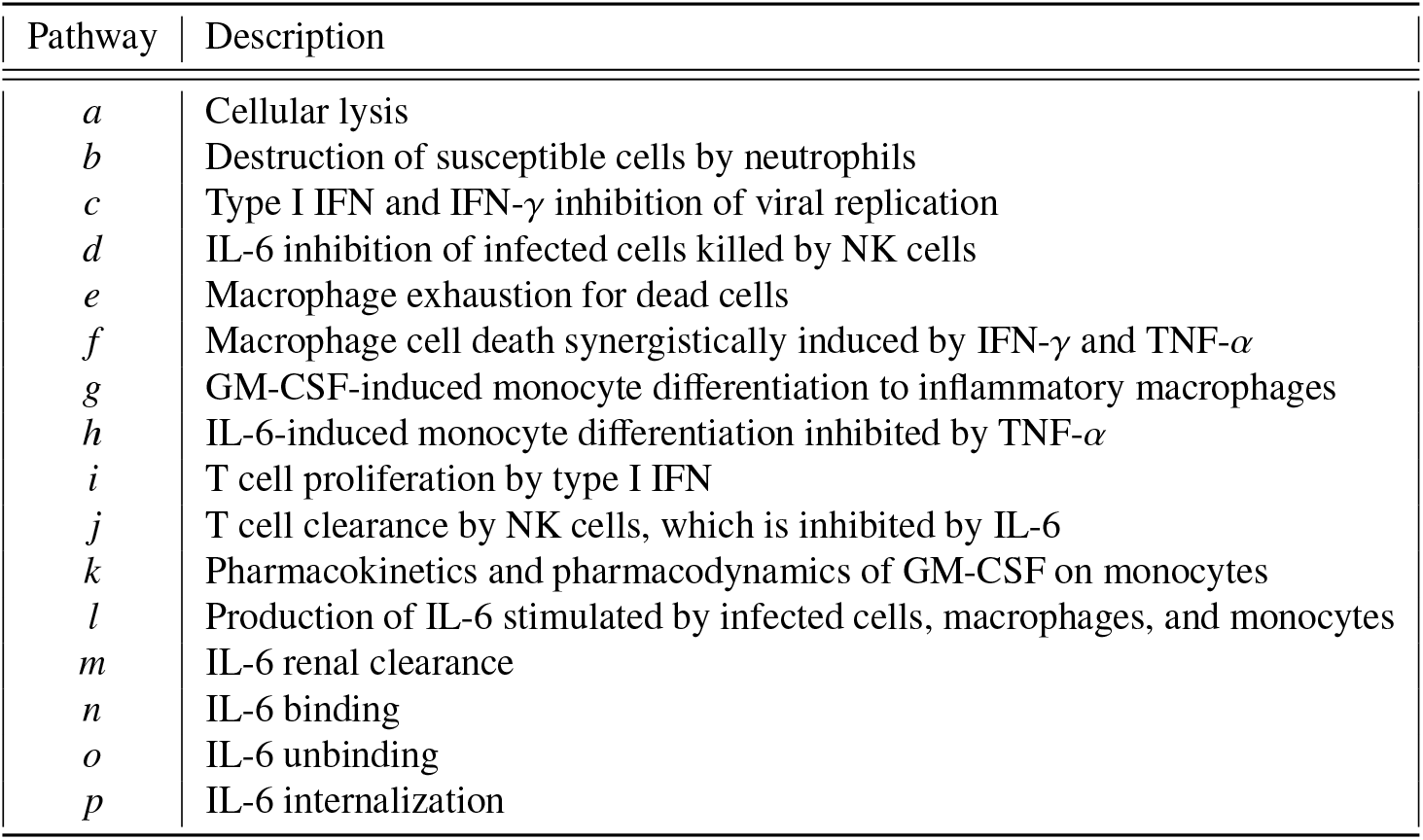
Description of selected biological pathways in Eqs. (1)–(23).

| Pathway | Description |
| --- | --- |
| $a$ | Cellular lysis |
| $b$ | Destruction of susceptible cells by neutrophils |
| $c$ | Type I IFN and IFN-γ inhibition of viral replication |
| $d$ | IL-6 inhibition of infected cells killed by NK cells |
| $e$ | Macrophage exhaustion for dead cells |
| $f$ | Macrophage cell death synergistically induced by IFN-γ and TNF-α |
| $g$ | GM-CSF-induced monocyte differentiation to inflammatory macrophages |
| $h$ | IL-6-induced monocyte differentiation inhibited by TNF-α |
| $i$ | T cell proliferation by type I IFN |
| $j$ | T cell clearance by NK cells, which is inhibited by IL-6 |
| $k$ | Pharmacokinetics and pharmacodynamics of GM-CSF on monocytes |
| $l$ | Production of IL-6 stimulated by infected cells, macrophages, and monocytes |
| $m$ | IL-6 renal clearance |
| $n$ | IL-6 binding |
| $o$ | IL-6 unbinding |
| $p$ | IL-6 internalization |

The complete list of model parameters and their values is summarized in tables in the Supplementary Materials. Throughout the model, interactions between cytokines and cells are described by a positive (stimulatory) Hill function:

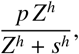

where *Z, s, h*, and *p* denote the interacting component, its half-effect value, the Hill coefficient, and the interaction rate, respectively [66, 67]. In what follows, the production (or recruitment/differentiation) rate of a given cell *X* by cytokine *Y* is denoted by *p*_*X,Y*_ ; the half-effect concentration (i.e., *s* in the Hill function above) of cytokine *X* acting on cell population *Y* by *ϵ*_*X,Y*_ ; and the half-effect concentration of cell *Y* affecting cytokine *X* by *η*_*X,Y*_. The natural death rate of cell *Y* is denoted by *d*_*Y*_, and the rate of induced death of cell *Y* by cell *Z* by *δ*_*Y,Z*_. Lastly, the proliferation rate of cell *Y* is denoted by *λ*_*Y*_, with carrying capacity *Y*_max_.

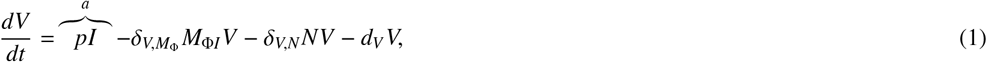

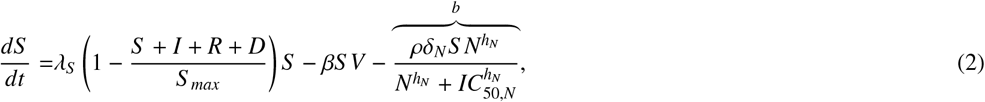

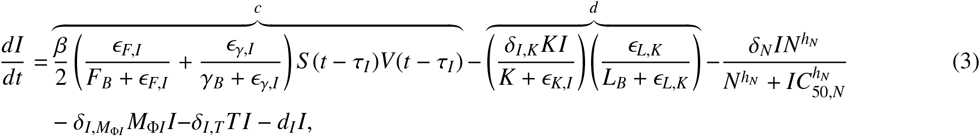

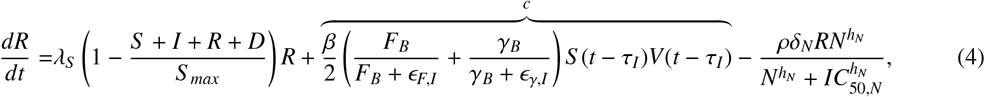

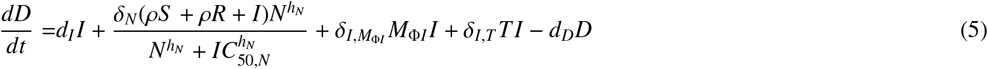

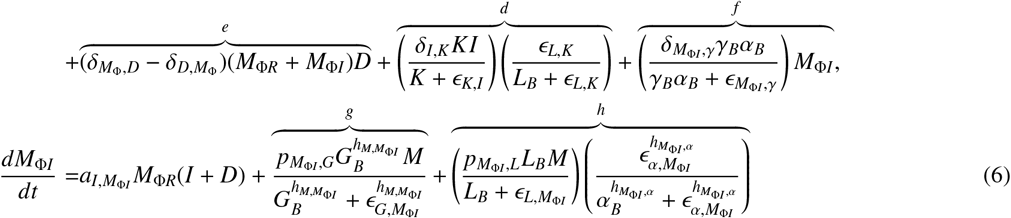

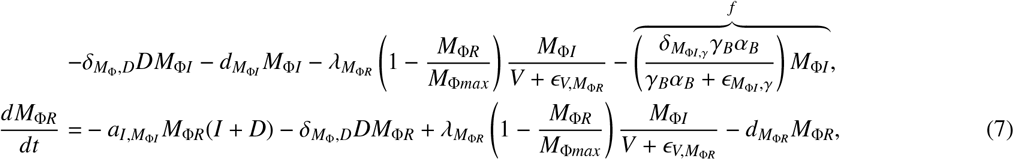

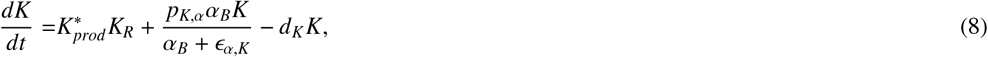

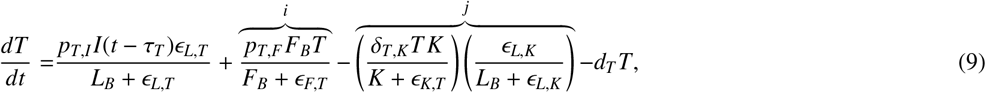

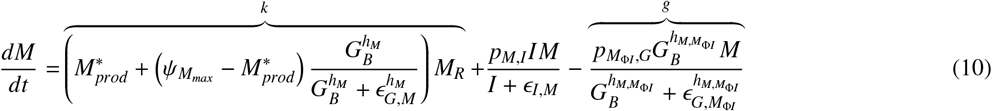

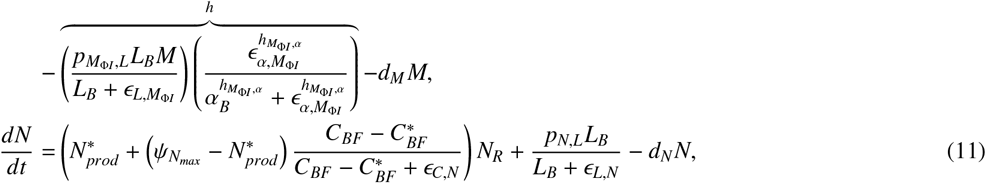

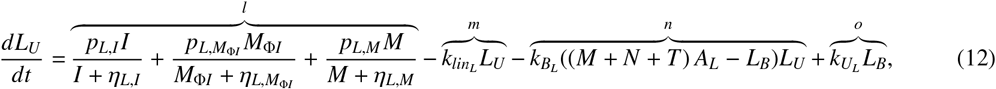

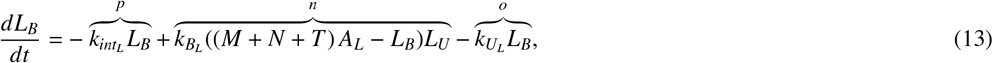

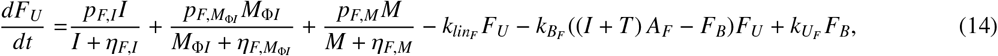

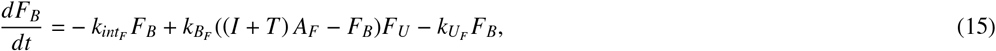

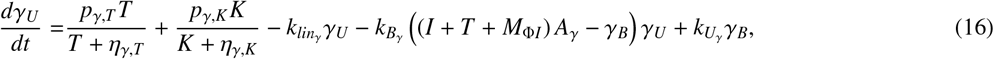

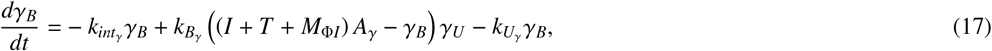

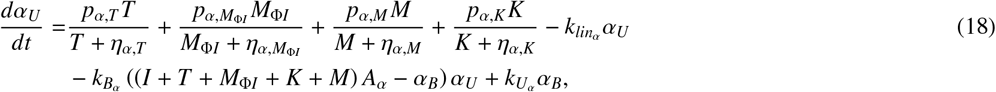

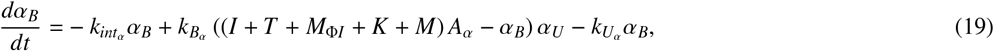

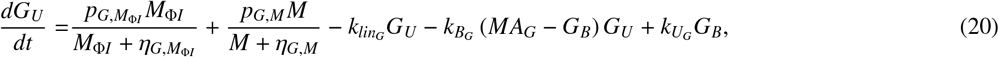

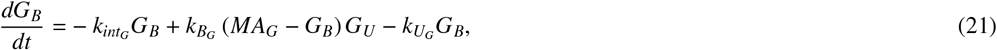

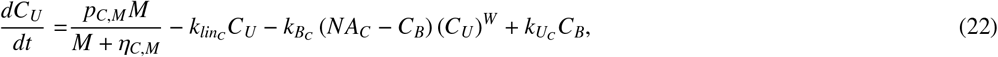

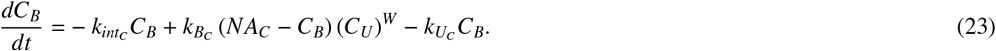

The dynamics for virus *V*, susceptible *S*, infected *I*, resistant *R*, and dead *D* cells (in Eqs. (1)–(5)) are adapted from [14] with the effect of additional terms including NK cells, IFN-*γ*, and TNF-*α* incorporated. Virus replicates at rate of *p* (pathway *a*) from infected cells while its clearance is modulated by inflammatory macrophages via apoptosis and neutrophils. The susceptible cell population is governed by a logistic growth at rate *λ*_*S*_, their infection interacting with virus at rate *β*, and bystander damage of epithelial cells by neutrophils (pathway *b*) through the release of reactive oxygen species [56, 57] using a Hill function.

The concentrations of bound type I IFN (*F*_*B*_) and bound IFN-*γ* (*γ*_*B*_) regulate the creation of infected and resistant cells [45, 68] in that the increased concentrations of both IFNs make more cells to become resistant to infection while making less cells infected, as reflected in Eqs. (3)–(4) or pathway *c*. The potency of this regulation is governed by the half-effect parameters, *ϵ*_*F,I*_ and *ϵ*_*γ,I*_. The delay *τ*_*I*_ represents the eclipse phase (i.e., time elapsed between successful cell infection and the start of virus production), after which infected cells begin to actively produce virus until undergoing their lytic death at rate *d*_*I*_. Though various immune cells contribute to the clearance of infected cells, in our model we consider only macrophages and CD8^+^ T cells, which lead to the apoptosis-induced death at rates 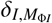 and *δ*_*I,T*_, respectively. Moreover, the last term of Eq. (3) (pathway *d*) shows that the killing of infected cells by NK cells is inhibited by IL-6 to include a biologically realistic representation of NK cell’s limited role in COVID-19 [69].

Dead cells (*D*) are accumulated through several ways: infected cell lysis *d*_*I*_, neutrophil damage of all epithelial cells *δ*_*N*_, the induced death of infected cells via macrophage phagocytosis 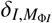, CD8^+^ T cell killing *δ*_*I,T*_, and NK killing *δ*_*I,K*_, macrophage exhaustion from clearing up dead cells 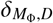 [70], and inflammatory macrophage death (of PANoptosis) synergistically modulated by both IFN-*γ* and TNF-*α* (see pathway *f*) with 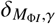 and half-effect 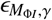 [46]. These dead cells degrade at a relatively high rate *d*_*D*_ [71] while being cleared up through phagocytosis by macrophages at rate 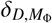.

The production of inflammatory macrophages (*M*_Φ*I*_) is regulated by three different pathways (acting individually or in concert): a transition from alveolar macrophages stimulated by infected and dead cells at a maximum rate 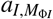, monocytes differentiation stimulated by GM-CSF (*G*_*U*_ and *G*_*B*_) with a maximum production rate 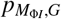 and half-effect 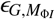 (in pathway *g*), and monocyte differentiation by IL-6 (*L*_*U*_ and *L*_*B*_) with a maximum production rate 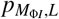 and half-effect 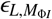 (in pathway *h*), as shown in the first three terms of Eq. (6). Specifically, the IL-6-dependent monocyte differentiation (i.e., pathway *h*) is inhibited by TNF-*α* because its presence derives monocyte differentiation more towards dendritic cells than macrophages [72, 73]. The next two terms in Eq. (6) represent the induced death of macrophage from clearing dead cells 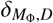 and natural death 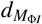, respectively. The logistic term with *M*_Φ*max*_ models that the inflammatory macrophages replenish the alveolar macrophage population in the lung as virus is cleared, thus, the replenishment of alveolar macrophages is inversely proportional to viral load with half-effect 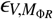. Resident macrophages are similarly modeled to undergo a switch to inflammatory macrophages stimulated by infected and dead cells, their replenishment as virus is cleared up, macrophage exhaustion from dead cell clearance at rate 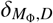, and natural death at rate 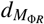.

NK cells (*K*) are modeled to stay at their homeostasis level with reservoir release rate 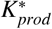 before infection, after which more NK cells are actively recruited by TNF-*α* [74] with a maximum rate *p*_*K,α*_ and half-effect *ϵ*_*α,K*_. They are cleared through natural death at rate *d*_*K*_.

The population of CD8^+^ T cells (*T*) is governed by the delay differential equation (9) where the first term describes cell recruitment at rate *p*_*T,I*_ with the delay *τ*_*T*_. The delay period accounts for the time needed for the arrival of effector CD8^+^ T cells at the infection site following dendritic cell activation, their migration to the lymph nodes, and the activation of CD8^+^ T cells. In this term, inhibition by IL-6 concentration (bound *L*_*B*_) is incorporated with half effect *ϵ*_*L,T*_ given that high concentrations of IL-6 result in CD8^+^ T cell exhaustion [75, 76]. In severe COVID-19 cases, impaired or delayed IFN signaling leads to defective virus-specific T cell activation and proliferation, which ultimately contributes to profound T cell lymphopenia [77, 78]. Thus, it is assumed that the expansion (or proliferation) process of the T cell population is mediated by type I IFN, as described in the pathway *i* with the Hill function. In addition, IL-6 has been found to contribute to NK cell dysfunction in target cell recognition [79] in severe COVID-19. Thus, T cells are modeled to be targeted (and cleared) by NK cells at rate of *δ*_*T,K*_ (pathway *j*) but this clearance is inhibited by the IL-6 concentration. T cell apoptosis occurs at rate *d*_*T*_.

Craig et al. [80] have shown that even under normal, healthy conditions (i.e., at homeostasis), endogeneous cy-tokine levels are not static or in a near-equilibrium state. To describe the pharmacokinetics and pharmacodynamics of cytokine binding and unbinding processes we use the framework established in [80]. The general form of the pharmacokinetic relationship is given by

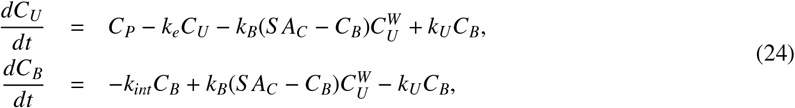

where *C*_*U*_ and *C*_*B*_ are unbound and bound cytokines with *k*_*U*_ and *k*_*B*_ being respective unbinding and binding rates, respectively. *C*_*P*_ is the rate of endogeneous cytokine production, *k*_*int*_ is the internalization rate of bound cytokine, and *k*_*e*_ is the elimination rate. Here, *W* is a stoichiometric constant and the sum concentration of all cells modulated by this cytokine *C* denoted by *S*. The calculation of a scaling factor *A*_*C*_ is determined by

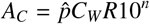

where 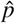 is a constant relating the stoichiometry between cytokine molecules and their receptors, *R* is the number of receptors to each cytokine on a cell’s surface and 10^*n*^ is a correcting factor for cellular units (see Eqs. (A.1)–(A.6)). The molecular weight is based on a standard calculation by dividing the cytokine’s molar mass (MM) by Avogadro’s number (i.e., *C*_*W*_ = *MM/*(6.02214 × 10^23^)). The cytokine concentration is then governed by the above expressions (24) and additional terms representing interactions of bound components with immune cells.

The dynamics for monocytes *M* and neutrophils *N* (Eqs. (10)–(11), respectively) interacting with cytokines are adapted from [14, 80], based on the cell-cytokine interaction using the Hill function, and the pharmacokinetics and pharmacodynamics of GM-CSF on monocytes and of G-CSF on neutrophils [81]. Specifically, as shown in pathway *k*, monocytes are recruited by bound GM-CSF [82] with bone marrow monocytes (*M*_*R*_) recruited at a homeostatic rate 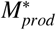. This recruitment rate increases up to maximum rate 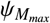 modulated by GM-CSF. In addition, infected cells produce monocytes at a maximal rate of *p*_*M,I*_ with half-effect *ϵ*_*I,M*_, but they will subsequently differentiate to inflammatory macrophages (as above in pathway *g*) or go through natural death at rate *d*_*M*_. Neutrophil recruitment from bone marrow reservoir neutrophils (*N*_*R*_) is similarly modeled with the bound fraction of G-CSF [80] (*C*_*BF*_ = *C*_*B*_(*t*)*/*(*A*_*C*_ *N*(*t*))) with its maximal value 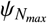. During the acute phase of inflammation, IL-6 is produced by endothelial cells, which leads to the attraction of neutrophils [83]. This recruitment is described in the second term of Eq. (11) with maximal rate *p*_*N,L*_ and half-effect parameter *ϵ*_*L,N*_. Neutrophils die at rate *d*_*N*_.

Model equations for cytokine production (Fig. 1B) are provided in Eqs. (12)–(23). Unbound IL-6 (*L*_*U*_) is produced by infected cells, inflammatory macrophages, and monocytes (pathway *l*), with bound IL-6 (*L*_*B*_) resulting from binding to receptors on the surface of neutrophils, CD8^+^ T cells and monocytes (pathway *n*). Though unbound type I IFNs (*F*_*U*_) are known to be secreted by multiple cell types upon viral infection, including lymphocytes, macrophages, endothelial cells and fibroblasts [68], we model its unbound production from infected cells, inflammatory macrophages, and monocytes, and its binding to receptors on both CD8+ T cells and infected cells to produce bound type I IFN (*F*_*B*_). While type I IFN is secreted upon viral infection, IFN-*γ* (*γ*_*U*_ and *γ*_*B*_) is secreted later due to delay in CD8^+^ T cell arrival at the infection site. Unbound IFN-*γ* production is modulated by CD8^+^ T cells and NK cells with bound INF-*γ* produced via binding to receptors of infected cells, CD8^+^ T cells, and macrophages. Unbound TNF-*α* (*α*_*U*_) are produced by inflammatory macrophages, monocytes, and NK and CD8^+^ T cells. Though the programmed death of infected cells induced by TNF-*α* [84] was not incorporated due to its insignificant effect in our model dynamics, binding of TNF-*α* to the receptors of infected as well as all other immune cells interacting with TNF-*α* is added in Eq. (18). Unbound GM-CSF (*G*_*U*_) is assumed to be produced from inflammatory macrophages and monocytes with bound GM-CSF (*G*_*B*_) produced via binding to monocyte receptors. The production of GM-CSF by CD8^+^ T cells [85] is excluded due to its insignificant effect in the full model’s dynamics. Lastly, unbound G-CSF (*C*_*U*_) is secreted by monocytes, and bind to neutrophil receptors to produce bound G-CSF (*C*_*B*_).

### 2.3 Parameter estimation and numerical simulation

Model parameters for our model (Eqs. (1)–(23)) are provided in Supplementary Materials, and they were determined through direct extraction from literature and curve fitting to capture empirical effects (by following similar data-fitting methods, e.g., `fmincon` or `lsqnonlin` functions in MATLAB) with data used in [14]. Initial values for the model variables are summarized in Table B.4. Data needed for NK cells, two added cytokines IFN-*γ* and TNF-*α*, and their interactions with other components in the model were adapted from either existing data for SARS-CoV-2, or previous work for MERS-CoV or SARS-CoV [86, 87], if needed and applicable. Any remaining parameters were calculated to preserve homeostatic equilibrium in healthy condition (i.e., in the absence of infection). A comprehensive description of parameter estimation process can be found in Supplementary Materials.

With the model with all calibrated parameters, we numerically solved our model using ode45 and ddesd in MATLAB to conduct the model validation against data and obtain all simulation results that follow. All codes used for model analysis are provided to replicate our results presented herein. See Data Availability for more information.

## 3. Main Results

### 3.1. Model predictions for mild versus severe disease

To identify key drivers that differentiate severe from mild dynamics, we numerically simulated our within-host model (Eqs. (1)–(23)) with two contrasting parameter sets: a baseline set to reproduce mild disease dynamics and an adjusted set to recapitulate severe dynamics. In clinical COVID-19 studies [63, 88, 89, 90], patients with severe disease were reported to have significantly lower levels of type I IFN, along with elevated monocyte and IL-6 concentrations. Elevated IL-6, in particular, correlates with NK-cell dysfunction due to its inhibitory effect on cytolytic activity of NK cells. In addition, IFN-*γ* production by CD8^+^ T cells is impaired because the ratio of CD4^+^ to CD8^+^ T cells increases, and CD8^+^ T cells produce more IFN-*γ* than CD4^+^ T cells [91].

To generate severe dynamics based on these observations, the rates of type I IFN production by infected cells (*p*_*F,I*_), IFN-*γ* production by T cells (*p*_*γ,T*_), and the half-effect parameter for IL-6 inhibition of NK-cell cytotoxicity (*ϵ*_*L,K*_) were decreased, whereas the production rate of monocytes by infected cells (*p*_*M,I*_) and the half-effect coefficient for type I IFN production by inflammatory macrophages 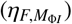 were increased. Model predictions are compared in Fig. 2, with mild disease shown as black solid lines and severe disease as red dashed lines. For model validation, viral loads from SARS-CoV-2 infection in humans reported in [92] (based on data from Singapore [93] and Germany [94]) were used, as indicated by open circles in Fig. 2A.

**Figure 2.**
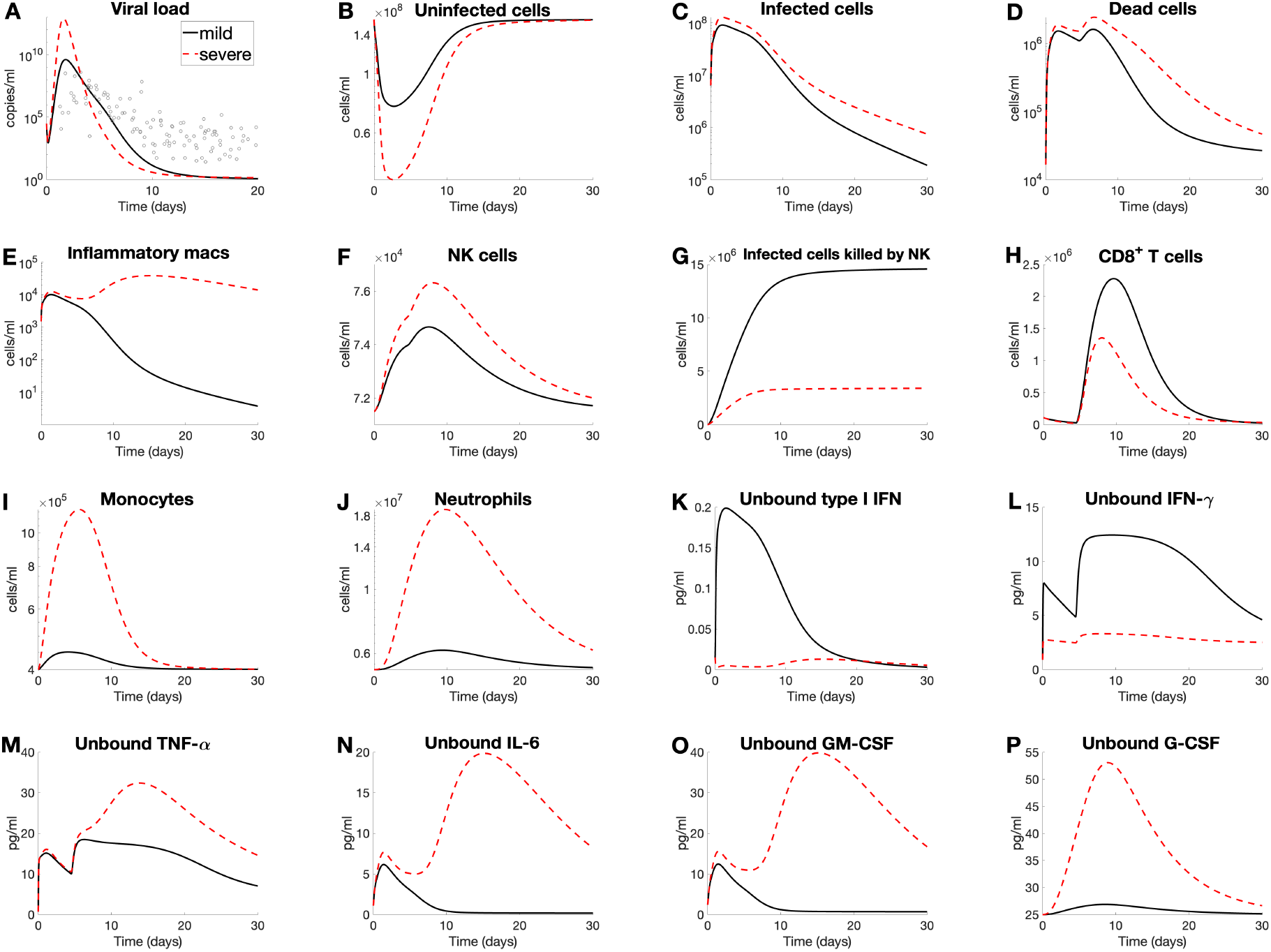
Model predictions for mild versus severe COVID-19 dynamics. Mild disease (solid lines) dynamics obtained by solving Eqs. (1)–(23) with baseline parameters summarized in Supplementary Materials. Viral load data (open circles) from eight human patients (three from Singapore and five from Germany by Goyal et al. [92]) are overlayed with predicted viral dynamics. Severe disease case (dashed lines) obtained by modifying selected parameters; decreasing production rate of type I IFN (via *p*_*F,I*_ and 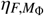), increasing monocyte recruitment rate (*p*_*M,I*_), decreasing NK cell’s cytotoxicity (*ϵ*_*L,K*_), and decreasing production rate of IFN-*γ* by T cells (*p*_*γ,T*_). The list of severe parameter values is summarized in Table B.5. The graphs depict the predicted concentrations of individual cell types or cytokines over the course of a 30-day infection. Only unbound cytokine concentrations (i.e., panels **K**–**P**) are shown, as these are clinically measurable and represent the pool available for receptor binding and downstream signaling. Time-series predictions of all other variables (i.e., including resident macrophages and all bound cytokines) are provided in Supplementary Materials.

The mild case shows effective clearance of the virus within ∼ 10 days, with all cell populations and cytokine concentrations returning to their respective homeostatic levels and only minimal tissue damage (Fig. 2B). Specifically, the innate immune response is restrained: inflammatory macrophages, NK cells, monocytes, and neutrophils reach their peaks within 10 days post-infection (Figs. 2E–F and 2I–J), followed by CD8^+^ T-cell activation peaking after day 10 (Fig. 2H). The cytokine dynamics in mild disease demonstrate a rapid response in type I IFN, IL-6, and GM-CSF (Figs. 2K, 2N, and 2O, respectively). The dynamics of IFN-*γ* and TNF-*α* (Figs. 2L–M) are distinct in that an initial rise occurs immediately after infection, followed by a rapid decline, after which concentrations reach their highest peak at around one week and remain at a relatively stable level for another week.

Using the adjusted parameter set, the model exhibits a dramatic shift in disease progression. Although the predicted viral load (Fig. 2A) remains comparable to that of mild infection, the severe scenario is characterized by a cytokine storm, with heightened levels of TNF-*α*, IL-6, GM-CSF, and G-CSF, consistent with experimental findings [6, 7, 39, 95]. This is accompanied by elevated ratios of innate to adaptive immune cells (Figs. 2E–J) and substantial loss of healthy lung tissue (Fig. 2B).

Moreover, in severe cases the interferon response (Figs. 2K–L) is characterized by a delayed and weakened peak of type I IFNs and by a comparatively reduced IFN-*γ* peak relative to mild cases, aligning well with data presented in [6, 29, 90]. In contrast to mild disease, inflammatory macrophages and neutrophils remain elevated for at least 30 days after initial infection (Figs. 2E and 2J). Notably, inflammation remains high in severe disease, with increased TNF-*α* and GM-CSF concentrations, despite the virus being cleared slightly faster (∼1 day) than in mild disease (Fig. 2A). The peak of inflammatory macrophages rises from ∼10^4^ cells/ml (mild) to ∼10^6^ cells/ml (severe).

The model also reproduces the substantial reduction in CD8^+^ T-cell concentrations in severe case (Fig. 2H), indicative of T-cell lymphopenia, consistent with clinical observations in patients with severe COVID-19 [86, 96]. Interestingly, even though NK-cell concentrations increase (Fig. 2F), their cytotoxic ability against infected cells shows a marked threefold reduction (Fig. 2G). This indicates impaired functional capacity, leading to NK-cell dysfunction [97]. Despite varying only five parameters 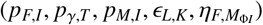, the model generates a distinct severe case, with cytokine dynamics qualitatively consistent with clinical observations for type I IFN [90, 98], IFN-*γ* [99], TNF-*α* [38], IL-6 [100, 101], and G-CSF [102, 103].

### 3.2. Sensitivity analysis identifies key pathways in the immune response to SARS-CoV-2 infection

To further examine the effects of parameter variation, we conducted a local sensitivity analysis (LSA) by increasing each parameter value by 5% from its baseline set (used for the mild dynamics in Fig. 2) and quantifying the resulting changes in selected variables to assess each parameter’s impact on host–pathogen dynamics. For each perturbation, we recorded predictions for key outputs over a 30-day window, including viral load, immune cell activation, tissue damage, and cytokine exposure. These outputs were then compared with baseline simulation results, and relative changes were calculated.

To facilitate visualization, model outputs were normalized using *z*-scores [104]. This standardization subtracts the mean and divides by the standard deviation, placing variables with different units on a common scale and enabling direct comparison of sensitivity magnitudes. Each parameter was assigned a sensitivity score, defined as the maximum absolute *z*-score across all outputs, and parameters were ranked accordingly; the top 23 most influential were selected for visualization in the heatmap of Fig. 3A. This shows the effects of perturbations on maximum viral load, maximum dead cells, minimum uninfected cells (tissue), peak immune cell concentrations (inflammatory macrophages, CD8^+^ T cells, and NK cells), maximum infected cells killed by NK cells, and maximum unbound cytokine levels (IL-6, type I IFN, IFN-*γ*, and TNF-*α*). Each heatmap cell represents the *z*-score–normalized deviation of a model output in response to a 5% parameter perturbation. The color scale ranges from blue (strong negative effect), through white (no effect), to red (strong positive effect), conveying both the magnitude and direction of influence while highlighting the most sensitive parameters. In addition, time-series dynamics for viral load, uninfected cells (tissue), infected cells killed by NK cells, and unbound IL-6, IFN-*γ*, and TNF-*α* concentrations after 5% parameter increases are shown in Figs. 3B–G. Finally, the maximum increase and maximum decrease values for each metric are listed in Table 3.

**Table 3:**
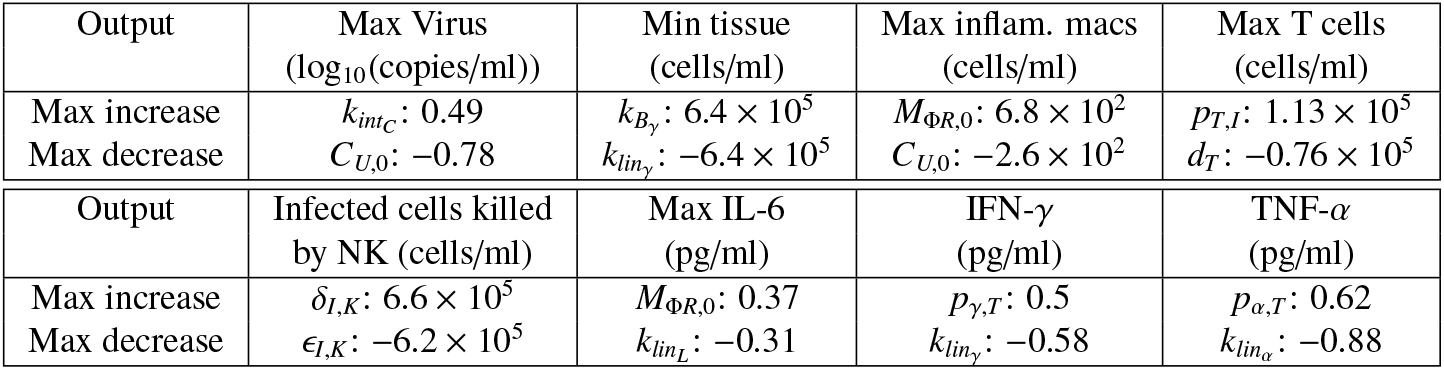
List of values for the maximum increase and maximum decrease of each metric.

**Figure 3.**
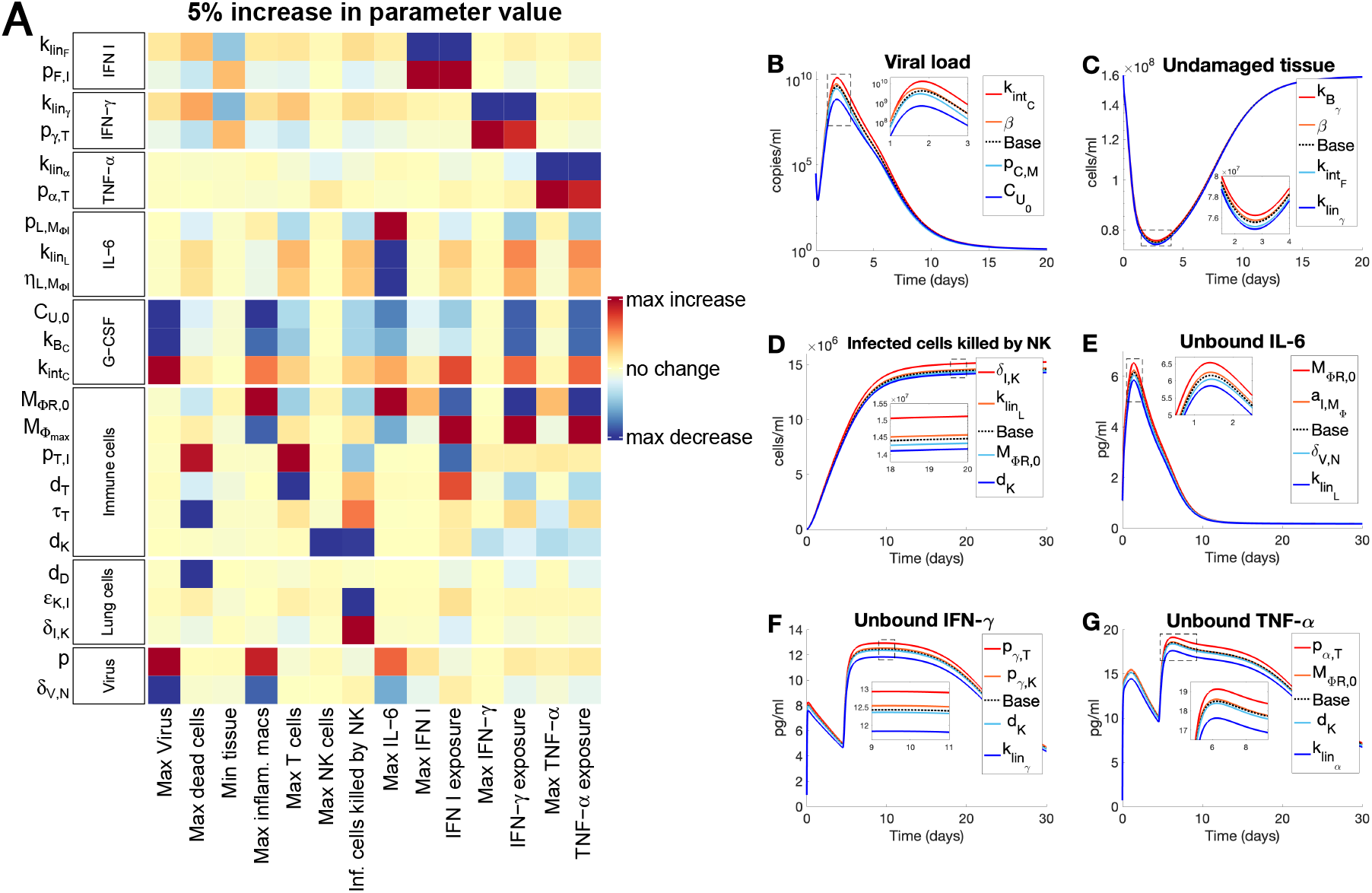
Top 23 sensitive parameters (with +5% perturbation) identified by local sensitivity analysis. Each parameter was increased by 5% from its originally estimated value (i.e., mild disease parameters in Fig. 2) and the resulting model dynamics were simulated. Predictions were then compared to baseline specifically for maximum viral load (max(*V*)), maximum concentration of dead cells (max(*D*)), minimum uninfected cells or tissues (min(*S* +*R*)), maximum concentrations of three immune cells–inflammatory macrophages (max(*M*_Ф*I*_)), CD8^+^ T cells (max(*T*)), and maximum concentration of NK cells (max(*K*)), and maximum concentration of infected cells killed by NK. The maximum concentrations of the four cytokines, IL-6 (max(*L*_*U*_)), type I IFN (max(*F*_*U*_)), IFN-*γ* (max(*γ*_*U*_)), and TNF-*α* (max(*α*_*U*_)) along with the corresponding total exposure to the last three cytokines. A) Heat map shows the magnitude of the change of each metric from a 5% increase in the parameter value compared to baseline, where blue and red represent the maximum and minimum values, respectively, observed in the output metric. The top 23 most sensitive parameters (categorized by a group of IFN I, IFN-*γ*, TNF-*α*, G-CSF, IL-6, immune cells, lung cells, and virus) are shown here, and for the complete parameter sensitivity results, see the Supplementary Materials. The explicit value of the maximum increase and maximum decrease of each metric is given in Table 3 below. The *z*-score normalization was applied to account for differences in scale and units across outputs, allowing for meaningful comparison of sensitivity magnitudes across all variables. **B)**–**G)**: Time-series dynamics of viral load, undamaged tissue (uninfected cells), infected cells killed by NK cells, and unbound IL-6, IFN-*γ*, and TNF-*α* given 5% increases in the noted parameters for each color. Colors of the curves correspond to the coloring of the heatmap in **A)**.

In Fig. 3A, a higher production rate of type I IFN by infected cells (*p*_*F,I*_) results in increased levels of unbound type I IFN, i.e., maximum level as well as the total exposure, based on max(*F*_*U*_) and total(*F*_*U*_) defined as the area under the unbound type I IFN curve over the course of exposure, respectively. This is accompanied by a moderate increase in undamaged tissue (min(*S* +*R*)) and a reduction in maximum dead cells (max(*D*)). These findings are consistent with the important role of type I IFN in promoting antiviral responses during early infection [105]. In contrast, increasing the renal clearance rate of unbound type I IFN 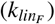 strongly decreases type I IFN levels, thereby impairing antiviral defense. This reduction is accompanied by moderate declines in uninfected and resistant cell populations (as also shown in Fig. 3C), while effects on immune cells and other cytokine levels remain modest. These results underscore the importance of sustained IFN signaling for viral control and are consistent with clinical observations of augmented clearance phenomena [106]. Sensitivity results for IFN-*γ* show similar trends: increasing the production rate of IFN-*γ* by T cells (*p*_*γ,T*_) promotes antiviral control, whereas increasing its renal clearance rate 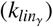 impairs it, as further supported by Fig. 3F.

Increasing the recruitment rate of T cells in response to infected cells (*p*_*T,I*_) enhances T-cell expansion (max(*T*)) and dead-cell accumulation, while reducing total type I IFN exposure. Minimum undamaged tissue, however, is unaffected. This suggests that enhanced T-cell recruitment promotes clearance of infected cells but has minimal impact on collateral tissue damage. Notably, NK-cell activity appears reduced despite stable NK-cell numbers (max(*K*)), due to the substantial increase in dead cells driven by infected-cell clearance [107, 108].

In addition, while increasing the initial number of resident macrophages (*M*_Φ*R*,0_) significantly decreases the exposure to type I IFN or IFN-*γ*, it leads to higher macrophage expansion (max(*M*_Φ*I*_)), elevated IL-6 levels (max(*L*_*U*_)), and increased maximum TNF-*α* levels (max(*α*_*U*_), shown in Fig. 3G). The increase IL-6 level slightly reduces the number of infected cells cleared by NK cells due to stronger IL-6–mediated inhibition of NK cytotoxicity [49], as can be seen by comparing Figs. 3D–E (especially in response to increased *M*_Φ*R*,0_). This reflects the strong role of macrophages in IL-6 production and its positive feedback on macrophage expansion [54]. While early macrophage presence and activation can enhance innate responses, excessive IL-6 promotes systemic inflammation and impairs balanced immunity, in part due to NK-cell dysfunction [109]. Moreover, the higher death rate of NK cells (*d*_*K*_) significantly reduces NK cell population, which consequently impairs their cytotoxic function (Fig. 3D) and decreases the production of IFN-*γ* and TNF-*α* (Figs. 3F–G).

A higher initial concentration of unbound G-CSF (*C*_*U*,0_) or its binding rate 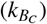 reduces peak viral load (Fig. 3B), dead cells, macrophages, T cells, and concentrations of all cytokines considered in our LSA. This suggests that early G-CSF availability facilitates effective infection control, limiting the need for excessive immune activation and reducing tissue damage. It also implies the therapeutic potential of early G-CSF supplementation, although careful consideration of timing and dosage would be critical to avoid overstimulation or suppression of cytokine-mediated immune dynamics following intervention [110].

Although not explicitly shown in the heatmap in Fig. 3A (but shown in the complete LSA results, provided in Supplementary Materials), a higher viral infection rate (*β*) markedly increases viral load, tissue damage, and immune dysregulation, as expected. Elevated *β* also drives higher levels of unbound IL-6 and type I IFN, consistent with uncontrolled viral replication leading to hyperinflammation and cytokine storm, as observed in patients with severe COVID-19 [111]. See Supplementary Materials for more information.

As noted in [14], the viral infectivity rate (*β*) can be influenced by viral mutations or varying densities of cellular ACE2 receptors between individuals. To assess whether the relative homogeneity of the immune response is preserved especially with new cytokine interactions added to our model, we varied the viral infectivity rate (*β*) by the same amount (0–50% with increments of 10%) and the corresponding results for viral load, inflammatory macrophages, T cells, NK cells, infected cells killed by NK, and unbound INF-*γ* and TNF-*α* were recorded in Figure 4. Although some variability is observed, both qualitative and quantitative behaviors including the shape and maximum concentrations of cytokines, including INF-*γ* and TNF-*α*, remain largely similar to the baseline case simulations. In addition, changes in inflammatory macrophages, T cells, NK cells, and NK cell cytotoxicity are minimal with their qualitative behaviors preserved. Importantly, viral load remains relatively the same except the higher peaks as *β* increases, suggesting that the increased infectivity alone is not sufficient to significantly alter disease progression and severity.

**Figure 4.**
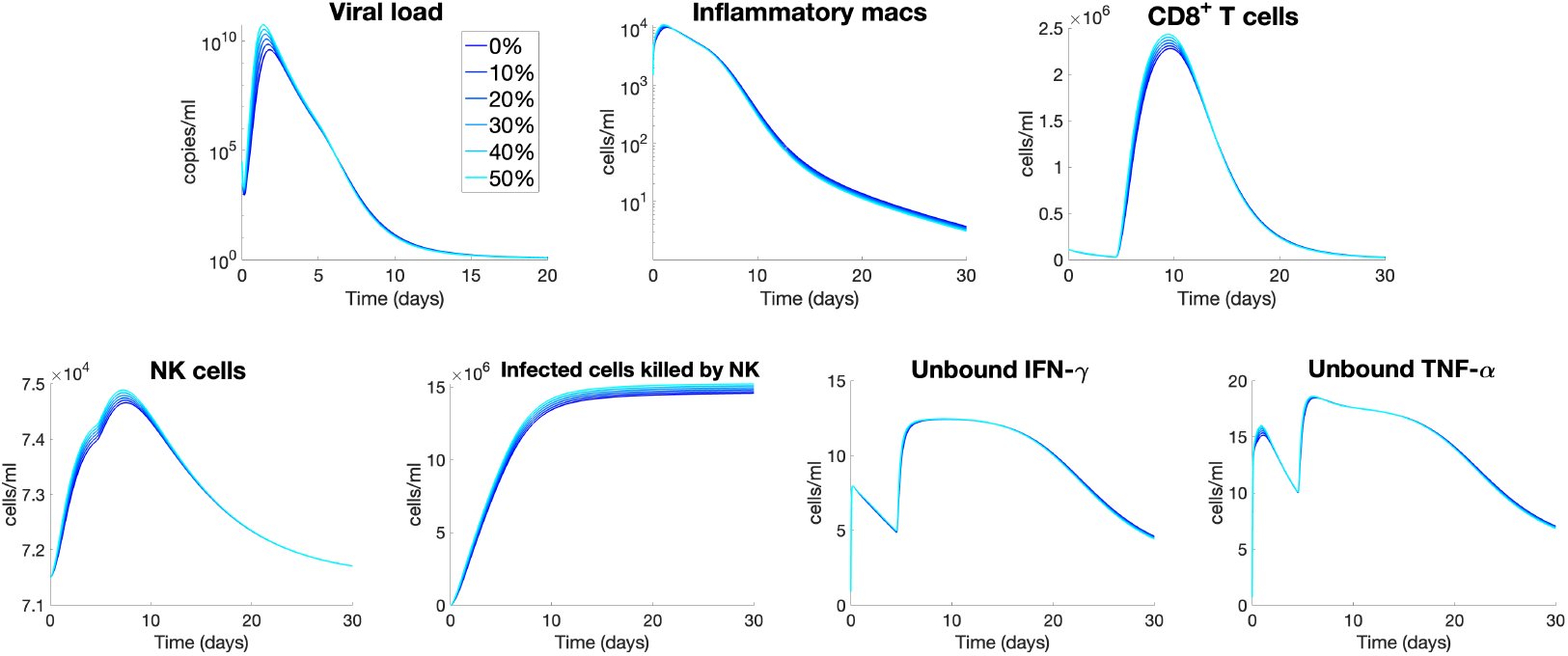
Viral infectivity rate *β* has small effects on overall immune dynamics. Dynamics for viral load, inflammatory macrophage, CD8^+^ T Cell, NK cells, and infected cells killed by NK cells, unbound IFN-*γ*, and unbound TNF-*α* are obtained using the baseline parameters (used for mild disease in Fig. 2) except increasing the viral infection rate *β* (by 0, 10, 20, 30, 40, 50%).

### 3.3. In silico knockdown analyses reveal how immune status influences disease outcomes

To assess the specific contributions of innate immune subsets to disease progression, we performed a series of *in silico* knockdown experiments. We focused on the mild disease course (Fig. 2, black solid line), since severe cases already involve systemic dysregulation, making it difficult to isolate the effects of individual cell types. In each simulation, recruitment of a given immune cell population was set to zero while initial abundances and death rates were left unchanged. Specifically, neutrophil recruitment was set to *p*_*N,L*_ = 0; inflammatory macrophage creation and recruitment to 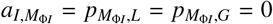 monocyte recruitment and differentiation to 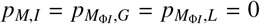 and NK cell recruitment to *p*_*K,α*_ = 0. The resulting dynamics are summarized in Fig. 5.

**Figure 5.**
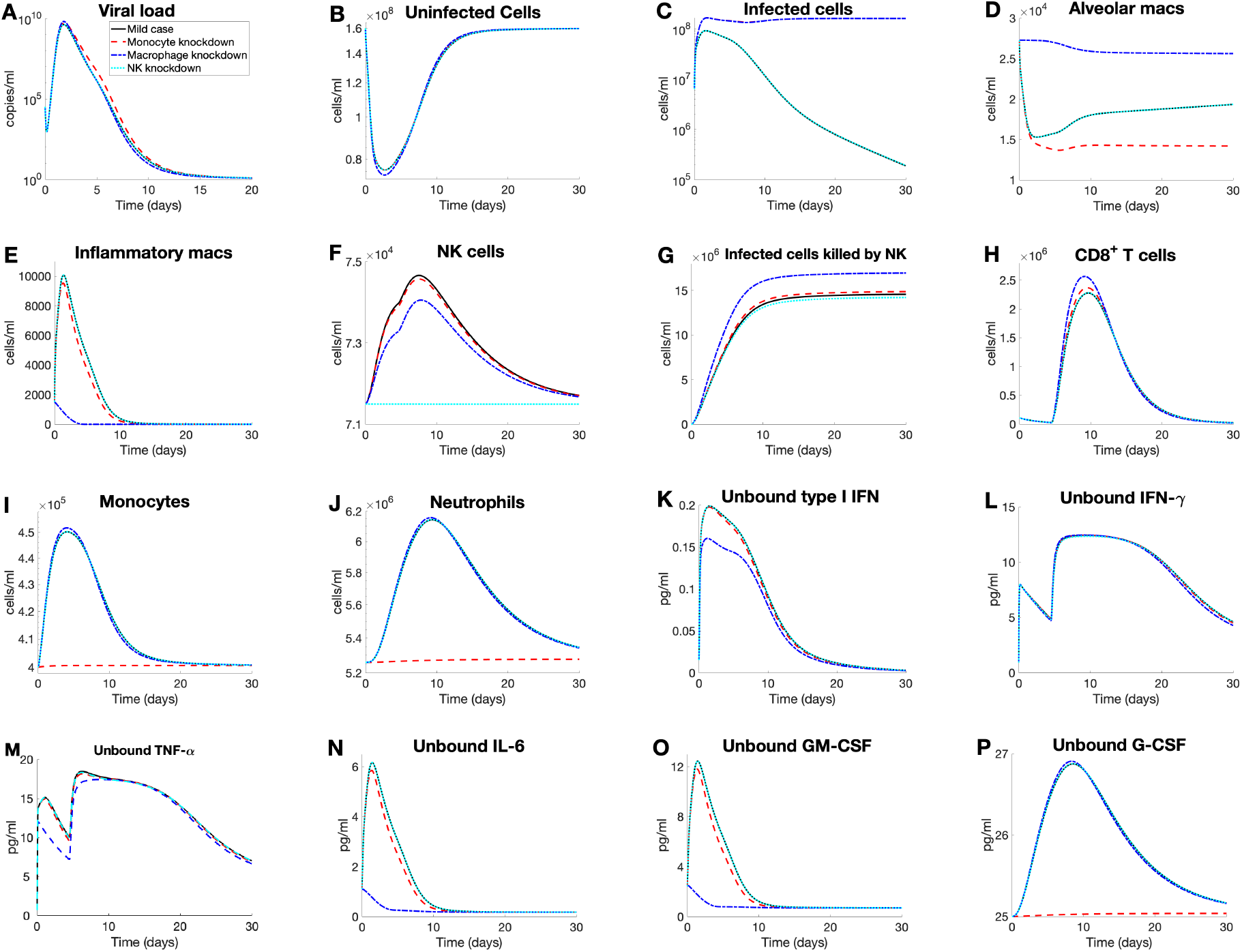
Effects of monocyte, macrophage, and NK cells knockdown on mild disease courses. *In silico* knockdown experiments in the mild disease scenario (as shown in Fig. 2; black solid line) were performed by implementing the recruitment knockdown of monocytes (i.e., no monocyte recruitment; red dash line), of inflammatory macrophages (i.e., not inflammatory macrophage creation via antigen stimulation or monocyte differentiation; blue dash-dot line), and of neutrophils (i.e., no neutrophil recruitment; cyan dot line), while keeping homeostasis and death rates the same. Model dynamics of the *in silico* knockdown are shown for **A)** viral load, **B)** uninfected cells, **C)** infected cells, **D)–J)**: immune cells and infected killed by NK cells (panel **G**), and **K)–P)**: unbound cytokines.

In the monocyte knockdown (red dashed line), loss of monocyte recruitment disrupted alveolar macrophages, NK-cell activity, neutrophil counts, and G-CSF concentration (Figs. 5D–F, J, P). Because monocytes are precursors to macrophages [112], their absence caused a marked reduction in resident macrophages. Monocytes also promote production of most cytokines except IFN-*γ* (Fig. 1B); accordingly, cytokine levels declined, with G-CSF showing the sharpest decrease (Fig. 5P). Reduced G-CSF lowered neutrophil recruitment (Fig. 5J). Although monocytes do not directly affect NK-cell numbers, the higher pool of infected cells increased NK-mediated killing (Fig. 5G). Viral load was slightly prolonged (Fig. 5A), indicating delayed clearance [113].

When inflammatory macrophages were knocked down (blue dash-dotted line), IL-6 and GM-CSF levels were most affected (Figs. 5N–O). Without macrophages, innate cytokine signaling was broadly impaired, reducing type I IFN, TNF-*α*, IL-6, and GM-CSF production (Figs. 5K, M–O), consistent with their central role in cytokine secretion (Fig. 1C). Viral control was also compromised, with sustained high numbers of infected cells (Fig. 5C) [114, 115]. Macrophage loss decreased TNF-*α* (Fig. 5M), reducing NK recruitment (Fig. 5F). However, the elevated infected-cell burden boosted NK-mediated killing (Fig. 5G), underscoring compensatory interactions among innate subsets.

In the NK-cell knockdown (cyan dotted line), cytokine dynamics were largely unchanged, but infected-cell clearance was modestly impaired (Fig. 5G) [116]. Viral load and uninfected-cell trajectories were nearly identical across knockdowns (Figs. 5A–C), illustrating the immune system’s ability to buffer the loss of individual components.

Across all knockdown scenarios, CD8^+^ T cells were maintained or slightly elevated relative to the mild case (Fig. 5H). This suggests that adaptive responses compensated for innate dysfunction, preserving viral clearance and IFN-*γ* dynamics (Figs. 5A,L). Importantly, none of the knockdowns reproduced the hallmarks of severe disease (e.g., loss of uninfected lung tissue, expansion of inflammatory macrophages; Figs. 5B,E). Thus, while individual innate-cell perturbations altered immune trajectories, severe disease emerges from systemic dysregulation rather than the loss of a single cell subset.

### 3.4 In silico cytokine knockdown investigations suggest therapeutic treatments for severe disease

To evaluate whether limiting specific pro-inflammatory cytokines can mitigate disease severity and point toward therapeutic strategies, we simulated knockdowns of IL-6, TNF-*α*, and IFN-*γ* on the severe disease trajectory (from Fig. 2, red dashed line), as summarized in Figure 6. In each case, cytokine production terms were set to zero for their modeled sources: for IL-6 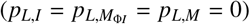, TNF-*α* (*p*_*α,T*_ = *p*_*α,M*_ = *p*_*α,K*_ = 0), and IFN-*γ* (*p*_*γ,K*_ = *p*_*γ,T*_ = 0). Note that though TNF-*α* is predominantly recruited by inflammatory macrophages, their recruitment rate (i.e., 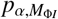) was kept at nonzero to mimic the most realistic medical knockdown intervention while focusing more on the potential effects of cytokine knockdown on NK cell function.

**Figure 6.**
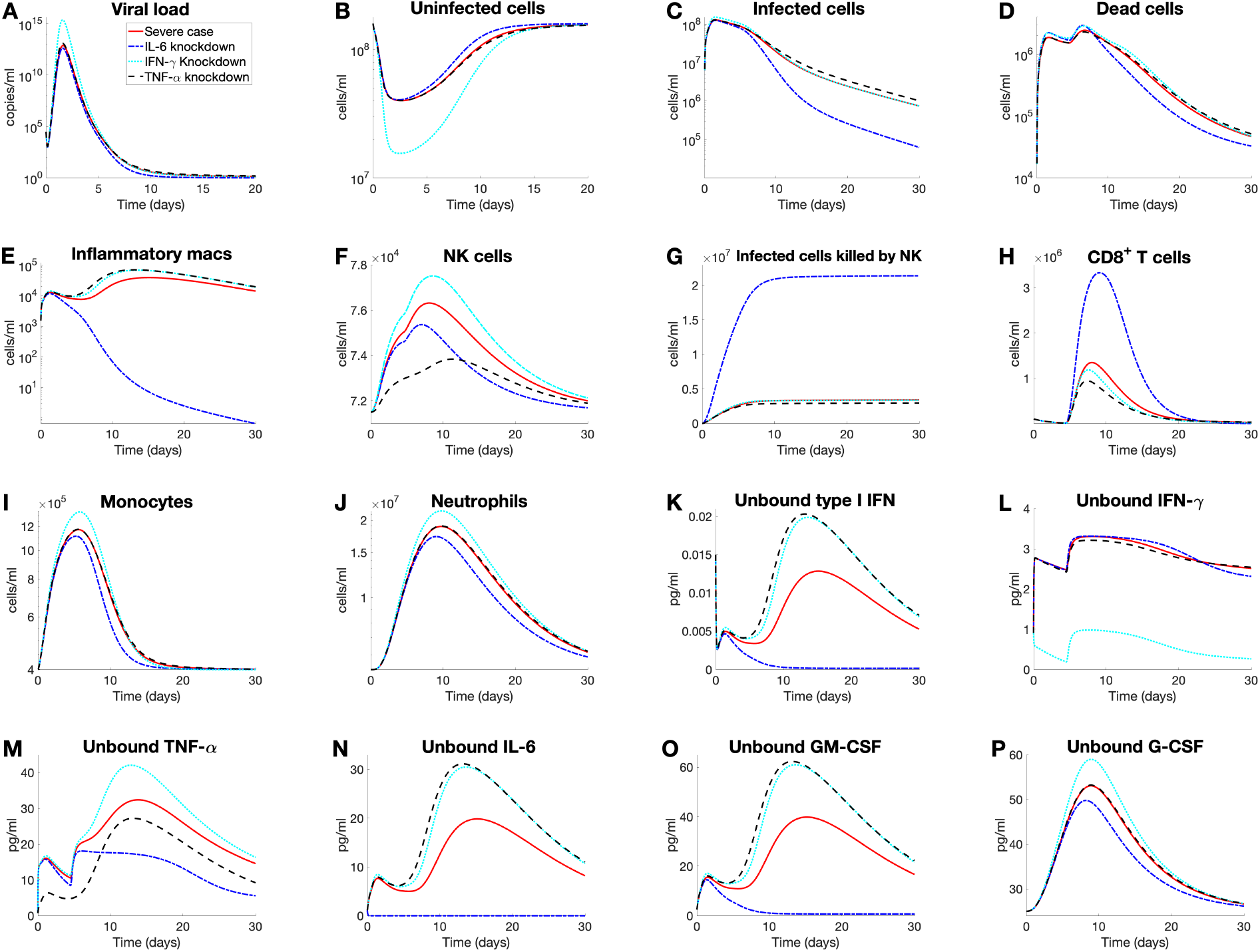
Effects of IL-6, TNF-*α*, and IFN-*γ* knockdown on severe disease courses. We performed *in silico* knockdown experiments in the severe disease scenario (Fig. 2; red line) by knocking down the production of IL-6 (by infected cells, inflammatory macrophages, and monocytes; blue dash-dot line), of TNF-*α* (by CD8^+^ T cells and monocytes; black dash line), and of INF-*γ* (by NK and CD8^+^ T cells; cyan dot line). Model dynamics of the *in silico* knockdown are plotted for **A)** viral load, **B)–D)** lung cells, **E)–J)** immune cells and infected cells killed by NK cells (panel **G**), and **K)–P)** unbound cytokines.

#### IL-6 knockdown

IL-6 knockdown scenario results in a markedly rapid decline in infected cells, less tissue damage, and higher population of CD8^+^ T cells, compared to TNF-*α* and IFN-*γ* knockdowns (Figs. 6B–D, H). This is consistent with clinical finding that IL-6 contributes to hyper-inflammation and cytokine storm, which can exacerbate the severity of COVID-19, for example, with elevated infected cells and tissue damage but reduced T-cell activity [109, 117, 118]. In addition, limiting IL-6 production leads to substantially reduced macrophage population, highlighting its role in macrophage activation and recruitment in lung tissue [52]. Specifically, given that IL-6 induces the differentiation of the recruited monocytes to inflammatory macrophages [54], its blockage could mitigate the immune response from excessive inflammation (Fig. 6E). In contrast, the CD8^+^ T cell response is notably enhanced by IL-6 knockdown, given the inhibitory role of IL-6 in T cell expansion [119], leading to an elevated peak comparable to a mild case, as can be seen by comparing Fig. 2H (black solid line) and Fig. 6H (blue dash-dot line).

IL-6 knockdown markedly lowers NK-cell abundance with peaks fall to ∼75% of the severe-case level (Fig. 6F), suggesting that, although IL-6 contributes to NK dysfunction in severe inflammation, it supports NK-cell maintenance under inflammatory stress and, most importantly, restores the ability for NK cells to kill infected cells (Fig. 6G).

In addition to immune cell activities, IL-6 knockdown significantly reduces the concentrations of all cytokines except IFN-*γ*. Both type I IFN and GM-CSF are suppressed (Figs. 6K and 6O, respectively), consistent with their dependence on monocytes and inflammatory macrophages (Figs. 1B and 6E). While IL-6 knockdown dampens macrophage-driven inflammation, as mentioned above, it also weakens early antiviral defenses and may impair viral clearance [120]. Moreover, the near-absence of GM-CSF and, consequently, a reduction in recruited monocytes (Figs. 6O and 6I, respectively) indicate IL-6 as a major upstream driver in severe disease [121]; given GM-CSF’s roles in leukocyte recruitment and lung tissue support [82], its depletion in the absence of IL-6 risks delayed resolution and prolonged infection. Overall, IL-6 knockdown, while beneficial in reducing inflammatory damage, may suppress critical immune functions required for effective viral clearance and tissue homeostasis.

The concentration of G-CSF, a cytokine central to neutrophil production, maturation, and function [81], is lowered by IL-6 knockdown (Fig. 6P). While the reduced IL-6 production may alleviate hyper-inflammation in severe COVID-19, the accompanying G-CSF drop can impair neutrophil responses (Fig. 6J) and weaken host defense [122], underscoring a therapeutic trade-off between dampening inflammation and preserving immune competence. Also, the IL-6/TNF-*α* axis shows reciprocal control: IL-6 knockdown reduces TNF-*α* by limiting macrophage activation, whereas TNF-*α* knockdown elevates IL-6 by removing regulatory restraint (Figs. 6M–N; Fig. 1). Therapeutically, perturbing either node can shift the balance toward under- or over-production of the other, with implications for inflammation control and tissue protection [123, 124]. Lastly, IL-6 knockdown exerts time-dependent effects on IFN-*γ* (Fig. 6L) in that its level rises around 10–20 days after symptom onset but drops below severe case after 20 days, suggesting more prolonged inflammation.

#### IFN-γ knockdown

IFN-*γ* knockdown produces a higher NK-cell peak than the severe case (Fig. 6F), consistent with IFN-*γ*-mediated negative feedback on NK cells [125]. Despite of increased NK cell population, however, IFN-*γ* blockage leads to the increased peak of viral load but lowered T-cell level (Figs. 6A and H, respectively) while having excessive inflammation, as well demonstrated by elevated concentrations for all other cytokines. This finding implies that IFN-*γ* is detrimental to viral clearance and T cells level [126, 127], and its loss destabilizes immune regulation.

#### TNF-α knockdown

Finally, TNF-*α* knockdown yields a stronger net anti-inflammatory effect in NK cell proliferation (Fig. 6F), reflecting TNF-*α*’s critical role in NK activation, proliferation, and survival [74]. The reduced NK cell population which subsequently descreases IFN-*γ* recruitment (as shown in Figs. 1B and 6L). Despite the overall reduction in inflammatory signaling, however, inflammatory macrophages increase slightly (Fig. 6E) because TNF-*α* regulates macrophage activity and limits monocyte differentiation to inflammatory macrophages. Thus, its loss may lessen macrophage apoptosis and remove this restraint, allowing accumulation [128]. Consistent with increased macrophage numbers, GM-CSF rises during TNF-*α* knockdown (Fig. 6O), as GM-CSF production tracks monocyte/macrophage abundance and TNF-*α* normally inhibits their differentiation/activation [72]. The observed elevation of GM-CSF following TNF-*α* suppression may indicate a compensatory mechanism possibly aimed at sustaining macrophage function despite the loss of TNF-*α* signaling. TNF-*α* knockdown also elevates type I IFN (Fig. 6K), suggesting that TNF-*α* may exert a negative regulatory effect on IFN signaling [129], possibly functioning as a feedback mechanism to limit excessive immune activation.

#### Synthesis

Overall, IL-6 knockdown mitigates hyperinflammation and restores T cell and NK cytotoxic function despite lowering NK counts (as can be seen in Figs. 6F–G), but at the cost of impaired cytokine support (type I IFN, GM-CSF, and G-CSF). TNF-*α* knockdown reduces inflammation but destabilizes macrophage control and suppresses NK activity. IFN-*γ* knockdown worsens infection despite transient NK expansion. Together, these results suggest that therapeutic benefit lies not in complete suppression but in *targeted modulation*, particularly of IL-6 and TNF-*α*, to balance inflammation control with preservation of antiviral immunity, consistent with clinical findings [11, 38, 39, 130].

## 4. Conclusion

In this study, we present a comprehensive within-host model of SARS-CoV-2 infection that highlights the pivotal roles of NK cells and cytokines, particularly IFN-*γ* and TNF-*α*, in shaping disease trajectories. Unlike previous models, ours incorporates specific immune cells and cytokines implicated in immune dysfunction and severe COVID-19 outcomes [29, 131]. By simulating their interactions, our framework provides insights into pathways that contribute both to successful viral clearance and to excessive inflammatory responses associated with acute lung injury and mortality. To our knowledge, this is the first attempt to primarily investigate key pathways of immune cell dysfunction, especially in the context of NK cells. By combining simulations, sensitivity analyses, and knockdown experiments, the model highlights the critical role of NK-cell function, the nuanced effects of IL-6 blockade, and the context-dependent impact of IFN-*γ* and TNF-*α* signaling. These insights not only align with emerging clinical observations but also underscore the need for personalized and combinatorial therapeutic strategies.

Despite comparable NK-cell counts in mild and severe cases, severe COVID-19 is marked by functional NK-cell impairment, often driven by elevated IL-6 levels. Clinically, high IL-6 correlates with reduced granzyme A-expressing NK cells, reflecting suppressed cytotoxicity [88]. Our NK-cell knockdown simulations (Fig. 5) recapitulated this, showing minimal effects on cytokine dynamics but a marked impairment in viral clearance. These findings reinforce the essential role of NK cells in early infection control and align with reports [64, 132] linking NK dysfunction to prolonged viral persistence and disease severity. Moreover, cytokines such as IL-6 and TGF-*β* exacerbate NK impairment by downregulating adhesion and cytotoxic functions [69], suggesting that therapeutic strategies to restore NK function, including IL-6 blockade, could enhance viral clearance and improve clinical outcomes.

Clinical trials of IL-6 inhibitors have produced mixed results. The WHO REACT meta-analysis of over 10,000 patients found that IL-6 receptor antagonists, when combined with glucocorticoids, reduced 28-day mortality, particularly in patients on non-invasive ventilation [133]. However, benefits were inconsistent in mechanically ventilated patients, and efficacy was contingent on concurrent glucocorticoid use, suggesting a synergistic rather than standalone role. Other studies (e.g., [134]) question IL-6 as a primary target, noting that levels in COVID-19 often fall below thresholds associated with efficacy in other inflammatory diseases. Potential risks, including secondary infections due to immunosuppression, further complicate its utility. This suggests that IL-6 blockade alone is insufficient but may contribute to benefit in well-selected patients, particularly in combination with glucocorticoids or JAK inhibitors [130]. Our findings highlight the importance of precision-medicine approaches and the use of biomarkers (e.g., serostatus [135]) to identify subgroups most likely to respond.

We also modeled the reported synergy between IFN-*γ* and TNF-*α*, which drives PANoptosis (i.e., programmed cell death) in macrophages [46]. Our multiplicative representation yielded minimal synergistic effects, despite clinical observations of high TNF-*α*, moderate IFN-*γ*, and elevated inflammatory macrophages [28, 115]. This discrepancy suggests that co-targeting TNF-*α* and IFN-*γ* may not provide the anticipated therapeutic benefit, in line with clinical results in [46].

During severe COVID-19, IFN-*γ* levels are difficult to quantify, with studies reporting both elevated and diminished concentrations [7, 28, 29, 30]. This uncertainty poses a challenge for modeling, as assumptions must be made regarding which IFN-*γ* range best reflects severe disease. While our model cannot resolve exact concentrations, it illustrates how varying IFN-*γ* levels influence immune dynamics. TNF-*α* knockdown simulations in severe cases (Fig. 6) exacerbated T-cell lymphopenia, elevated TNF-*α* and IL-6, and worsened outcomes. These results suggest that IFN-*γ* inhibition would likely be harmful, particularly in patients already exhibiting low IFN-*γ* levels.

Conversely, clinical studies during the early pandemic highlighted potential therapeutic roles for IFN-*γ* supplementation [136, 137, 138]. IFN-*γ* treatment was reported to aid in the management of community-acquired infections and ventilator-associated pneumonia (VAP) in COVID-19 patients [137]. Interestingly, its efficacy appeared bidirectional: in one study, severely ill patients with immune deficiencies recovered following IFN-*γ* administration [138], while another reported recovery in patients with hyper-inflammatory responses complicated by VAP [137]. In line with these reports, our simulations suggest that increasing IFN-*γ* in mild cases promoted greater tissue protection and improved immune responses, indicating that moderate IFN-*γ* supplementation could be beneficial. Further research is needed to determine whether there is a threshold above which IFN-*γ* becomes pathogenic in severe COVID-19. Together, these findings suggest that IFN-*γ* may be a broadly safe yet context-sensitive therapeutic option across diverse immunoprofiles.

Despite the complexity of our model and the breadth of results, this study has several limitations that should be addressed in future work. First, macrophage polarization was not explicitly modeled, although the M1 phenotype is closely linked to cytokines included in our framework such as IFN-*γ*, TNF-*α*, and GM-CSF [139]. Future extensions could incorporate macrophage polarization and exogenous drivers such as IL-4, IL-10, IL-13, and TGF-*β*. Second, we modeled the innate immune response in unvaccinated individuals. Given widespread vaccination, future work should account for adaptive immunity, including B-cell and antibody dynamics, and examine how vaccination alters cytokine-driven pathways. Finally, we did not include TNF-*α*–induced epithelial apoptosis, which some studies suggest may be significant [38]. Including these processes could further refine predictions of disease severity and therapeutic response.

## Supporting information

Complete Sensitivity Analysis Results

## CRediT authorship contribution statement

### Pagnapech Ngoun

Conceptualization, Methodology, Validation, Formal analysis, Investigation, Software, Visualization, Writing - original draft, Writing - review & editing.

### Nicolas Alvarez

Conceptualization, Methodology, Validation, Formal analysis, Investigation, Software, Visualization, Writing - original draft, Writing - review & editing.

### Ayesh Awad

Conceptualization, Methodology, Validation, Investigation, Software, Visualization, Writing - review & editing.

### Hwayeon Ryu

Conceptualization, Methodology, Validation, Formal analysis, Investigation, Software, Visualization, Writing - original draft, Writing - review & editing, Project administration, Funding acquisition.

## Acknowledgment

We thank Murilo Ferreira Lopes for his contribution to the early stage of model formulation and parameter estimation in this study.

## Data Availability

Matlab codes for parameter estimation and numerical simulation including knockdown experiments, and Matlab and R codes for LSA analysis (shown in Sec. 3) are all available on the Github repository, whose link will be published upon the acceptance of this paper.

## Appendix A. Cytokine’s average receptor number

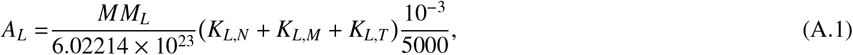

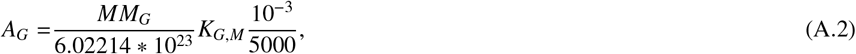

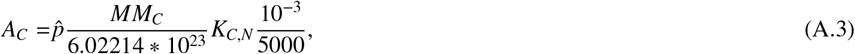

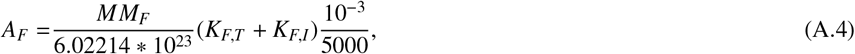

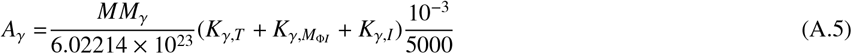

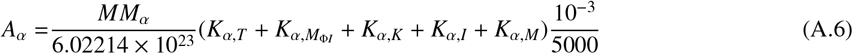

## Appendix B. Table of model parameters

The comprehensive list of model parameters and corresponding estimation process are provided in Supplementary Materials.

**Table B.4:** Initial conditions for model variables.

| Symbol | Description | Unit | Value | Reference |
| --- | --- | --- | --- | --- |
| $V_0$ | Initial viral load | $\log_{10}(\text{cop/ml})$ | 4.5 | [140] |
| $S_0$ | Initial susceptible cells | $10^9 \text{ cells/ml}$ | 0.16 | [141, 142] |
| $I_0$ | Initial infected cells | $10^9 \text{ cells/ml}$ | 0 | – |
| $R_0$ | Initial resistant cells | $10^9 \text{ cells/ml}$ | 0 | – |
| $M_{\Phi R,0}$ | Initial resident macrophages | $10^9 \text{ cells/ml}$ | $2.7 \times 10^{-5}$ | [141] |
| $M_{\Phi I,0}$ | Initial inflammatory macrophages | $10^9 \text{ cells/ml}$ | $1.5 \times 10^{-6}$ | H |
| $M_0$ | Initial monocytes | $10^9 \text{ cells/ml}$ | 0.0004 | [143] |
| $M_R$ | Initial reservoir monocytes | $10^9 \text{ cells/ml}$ | 0.0023 | [14] |
| $N_0$ | Initial neutrophils | $10^9 \text{ cells/ml}$ | 0.0053 | [80] |
| $N_R$ | Initial reservoir neutrophils | $10^9 \text{ cells/ml}$ | 0.0316 | [80] |
| $T_0$ | Initial $\text{CD8}^+$ T cells | $10^9 \text{ cells/ml}$ | $1.1 \times 10^{-4}$ | [144, 145] |
| $K_0$ | Initial NK cells | $10^9 \text{ cells/ml}$ | $7.15 \times 10^{-5}$ | [146] |
| $K_R$ | Initial reservoir NK cells | $10^9 \text{ cells/ml}$ | $1.1 \times 10^{-4}$ | [147] |
| $L_{U,0}$ | Initial unbound IL-6 | pg/ml | 1.1 | [148] |
| $L_{B,0}$ | Initial bound IL-6 | pg/ml | $1.4 \times 10^{-6}$ | H |
| $G_{U,0}$ | Initial unbound GM-CSF | pg/ml | 2.43 | [149] |
| $G_{B,0}$ | Initial bound GM-CSF | pg/ml | $1.6 \times 10^{-8}$ | H |
| $C_{U,0}$ | Initial unbound G-CSF | ng/ml | 0.025 | [80] |
| $C_{B,0}$ | Initial bound G-CSF | ng/ml | $6.5 \times 10^{-10}$ | [80] |
| $F_{U,0}$ | Initial unbound type I IFN | pg/ml | 0.015 | [150] |
| $F_{B,0}$ | Initial bound type I IFN | pg/ml | $1.1 \times 10^{-8}$ | H |
| $\gamma_{U,0}$ | Initial unbound IFN-γ | pg/ml | 0.91 | [151] |
| $\gamma_{B,0}$ | Initial bound IFN-γ | pg/ml | $1.432 \times 10^{-7}$ | H |
| $\alpha_{U,0}$ | Initial unbound TNF-α | pg/ml | 0.7 | [152] |
| $\alpha_{B,0}$ | Initial bound TNF-α | pg/ml | $1.3 \times 10^{-6}$ | H |

**Table B.5:** Parameter modifications for severe COVID-19 disease in Fig. 2 (H refers to homeostasis calculation).

| Symbol | Description | Unit | Mild Case | Severe Case | Reference |
| --- | --- | --- | --- | --- | --- |
| $p_{F,I}$ | Production of type I IFN by infected cells | pg/ml/day | 2.82 | 0.002 | [153] |
| $p_{M,I}$ | Monocyte recruitment rate by infected cells | 1/day | 0.22 | 1.2 | [154] |
| $\eta_{F,M\Phi I}$ | Cytokine production half-effect coefficient of type I IFN by inflammatory macrophages | $10^9 \text{ cells/ml}$ | $1.3 \times 10^{-6}$ | $2 \times 10^{-4}$ | H |
| $\epsilon_{L,K}$ | Half effect constant for IL-6 inhibition of NK cell cytotoxicity | pg/ml | $2.102 \times 10^{-5}$ | $2.102 \times 10^{-6}$ | [79] |
| $p_{\gamma,T}$ | Production rate of IFN-γ by T cells | pg/ml/day | 300 | 30 | [155] |

