## Supplementary Materials for "Immune Cell Dysfunction of SARS-CoV-2: Mathematical Modeling of the Within-Host Immune Dynamics"

---

---

### 1. Model Equations

$$\frac{dV}{dt} = \overbrace{pI}^{a} - \delta_{V,M_\Phi} M_{\Phi I} V - \delta_{V,N} N V - d_V V, \tag{1}$$

$$\frac{dS}{dt} = \lambda_S \left(1 - \frac{S + I + R + D}{S_{max}}\right) S - \beta S V - \overbrace{\frac{\rho \delta_N S N^{h_N}}{N^{h_N} + IC_{50,N}^{h_N}}}^{b}, \tag{2}$$

$$\begin{aligned}\frac{dI}{dt} =& \overbrace{\frac{\beta}{2}\left(\frac{\epsilon_{F,I}}{F_B + \epsilon_{F,I}} + \frac{\epsilon_{\gamma,I}}{\gamma_B + \epsilon_{\gamma,I}}\right) S(t-\tau_I) V(t-\tau_I)}^{c} - d_I I \\ & - \frac{\delta_N I N^{h_N}}{N^{h_N} + IC_{50,N}^{h_N}} - \delta_{I,M_{\Phi I}} M_{\Phi I} I - \delta_{I,T} T I - \overbrace{\left(\frac{\delta_{I,K} K I}{K + \epsilon_{K,I}}\right)\left(\frac{\epsilon_{L,K}}{L_B + \epsilon_{L,K}}\right)}^{d},\end{aligned} \tag{3}$$

$$\frac{dR}{dt} = \lambda_S \left(1 - \frac{S + I + R + D}{S_{max}}\right) R + \overbrace{\frac{\beta}{2}\left(\frac{F_B}{F_B + \epsilon_{F,I}} + \frac{\gamma_B}{\gamma_B + \epsilon_{\gamma,I}}\right) S(t-\tau_I) V(t-\tau_I)}^{c} - \frac{\rho \delta_N R N^{h_N}}{N^{h_N} + IC_{50,N}^{h_N}}, \tag{4}$$

$$\begin{aligned}\frac{dD}{dt} =& d_I I + \frac{\delta_N (\rho S + \rho R + I) N^{h_N}}{N^{h_N} + IC_{50,N}^{h_N}} + \delta_{I,M_{\Phi I}} M_{\Phi I} I + \delta_{I,T} T I - d_D D \\ & + \overbrace{(\delta_{M_\Phi,D} - \delta_{D,M_\Phi})(M_{\Phi R} + M_{\Phi I}) D}^{e} + \overbrace{\left(\frac{\delta_{I,K} K I}{K + \epsilon_{K,I}}\right)\left(\frac{\epsilon_{L,K}}{L_B + \epsilon_{L,K}}\right)}^{d} + \overbrace{\left(\frac{\delta_{M_{\Phi I},\gamma} \gamma_B \alpha_B}{\gamma_B \alpha_B + \epsilon_{M_{\Phi I},\gamma}}\right) M_{\Phi I}}^{f},\end{aligned} \tag{5}$$

$$\begin{aligned}\frac{dM_{\Phi I}}{dt} =& a_{I,M_{\Phi I}} M_{\Phi R} (I + D) + \overbrace{\frac{p_{M_{\Phi I},G} G_B^{h_{M,M_{\Phi I}}} M}{G_B^{h_{M,M_{\Phi I}}} + \epsilon_{G,M_{\Phi I}}^{h_{M,M_{\Phi I}}}}}^{g} + \overbrace{\left(\frac{p_{M_{\Phi I},L} L_B M}{L_B + \epsilon_{L,M_{\Phi I}}}\right)\left(\frac{\epsilon_{\alpha,M_{\Phi I}}^{h_{M_{\Phi I},\alpha}}}{\alpha_B^{h_{M_{\Phi I},\alpha}} + \epsilon_{\alpha,M_{\Phi I}}^{h_{M_{\Phi I},\alpha}}}\right)}^{h} \\ & - \delta_{M_\Phi,D} D M_{\Phi I} - d_{M_{\Phi I}} M_{\Phi I} - \left(1 - \frac{M_{\Phi R}}{M_{\Phi max}}\right) \frac{\lambda_{M_{\Phi R}} M_{\Phi I}}{V + \epsilon_{V,M_{\Phi R}}} - \overbrace{\left(\frac{\delta_{M_{\Phi I},\gamma} \gamma_B \alpha_B}{\gamma_B \alpha_B + \epsilon_{M_{\Phi I},\gamma}}\right) M_{\Phi I}}^{f},\end{aligned} \tag{6}$$

---

*Corresponding author

$$\frac{dM_{\Phi R}}{dt} = -a_{I,M_{\Phi I}} M_{\Phi R}(I+D) - \delta_{M_{\Phi},D} D M_{\Phi R} + \left(1 - \frac{M_{\Phi R}}{M_{\Phi max}}\right) \frac{\lambda_{M_{\Phi R}} M_{\Phi I}}{V + \epsilon_{V,M_{\Phi R}}} - d_{M_{\Phi R}} M_{\Phi R}, \tag{7}$$

$$\frac{dK}{dt} = K^*_{prod} K_R + \frac{p_{K,\alpha} \alpha_B K}{\alpha_B + \epsilon_{\alpha,K}} - d_K K, \tag{8}$$

$$\frac{dT}{dt} = \frac{p_{T,I} I(t-\tau_T)\epsilon_{L,T}}{L_B + \epsilon_{L,T}} + \overbrace{\frac{p_{T,F} F_B T}{F_B + \epsilon_{F,T}}}^{i} - \overbrace{\left(\frac{\delta_{T,K} T K}{K + \epsilon_{K,T}}\right)\left(\frac{\epsilon_{L,K}}{L_B + \epsilon_{L,K}}\right)}^{j} - d_T T, \tag{9}$$

$$\frac{dM}{dt} = \overbrace{\left(M^*_{prod} + \left(\psi_{M_{max}} - M^*_{prod}\right) \frac{G_B^{h_M}}{G_B^{h_M} + \epsilon_{G,M}^{h_M}}\right) M_R}^{k} + \frac{p_{M,I} I M}{I + \epsilon_{I,M}} - \overbrace{\frac{p_{M_{\Phi I},G} G_B^{h_{M,M_{\Phi I}}} M}{G_B^{h_{M,M_{\Phi I}}} + \epsilon_{G,M_{\Phi I}}^{h_{M,M_{\Phi I}}}}}^{g} \tag{10}$$

$$- \overbrace{\left(\frac{p_{M_{\Phi I},L} L_B M}{L_B + \epsilon_{L,M_{\Phi I}}}\right)\left(\frac{\epsilon_{\alpha,M_{\Phi I}}^{h_{M_{\Phi I},\alpha}}}{\alpha_B^{h_{M_{\Phi I},\alpha}} + \epsilon_{\alpha,M_{\Phi I}}^{h_{M_{\Phi I},\alpha}}}\right)}^{h} - d_M M,$$

$$\frac{dN}{dt} = \left(N^*_{prod} + \left(\psi_{N_{max}} - N^*_{prod}\right) \frac{C_{BF} - C^*_{BF}}{C_{BF} - C^*_{BF} + \epsilon_{C,N}}\right) N_R + \frac{p_{N,L} L_B}{L_B + \epsilon_{L,N}} - d_N N, \tag{11}$$

$$\frac{dL_U}{dt} = \overbrace{\frac{p_{L,I} I}{I + \eta_{L,I}} + \frac{p_{L,M_{\Phi I}} M_{\Phi I}}{M_{\Phi I} + \eta_{L,M_{\Phi I}}} + \frac{p_{L,M} M}{M + \eta_{L,M}}}^{l} - \overbrace{k_{lin_L} L_U}^{m} - \overbrace{k_{B_L}((M+N+T)A_L - L_B)L_U}^{n} + \overbrace{k_{U_L} L_B}^{o}, \tag{12}$$

$$\frac{dL_B}{dt} = -\overbrace{k_{int_L} L_B}^{p} + k_{B_L}((M+N+T)A_L - L_B)L_U - k_{U_L} L_B, \tag{13}$$

$$\frac{dF_U}{dt} = \frac{p_{F,I} I}{I + \eta_{F,I}} + \frac{p_{F,M_{\Phi I}} M_{\Phi I}}{M_{\Phi I} + \eta_{F,M_{\Phi I}}} + \frac{p_{F,M} M}{M + \eta_{F,M}} - k_{lin_F} F_U - k_{B_F}((I+T)A_F - F_B)F_U + k_{U_F} F_B, \tag{14}$$

$$\frac{dF_B}{dt} = -k_{int_F} F_B + k_{B_F}((I+T)A_F - F_B)F_U - k_{U_F} F_B, \tag{15}$$

$$\frac{d\gamma_U}{dt} = \frac{p_{\gamma,T} T}{T + \eta_{\gamma,T}} + \frac{p_{\gamma,K} K}{K + \eta_{\gamma,K}} - k_{lin_\gamma} \gamma_U - k_{B_\gamma} \left((I + T + M_{\Phi I}) A_\gamma - \gamma_B\right) \gamma_U + k_{U_\gamma} \gamma_B, \tag{16}$$

$$\frac{d\gamma_B}{dt} = -k_{int_\gamma} \gamma_B + k_{B_\gamma} \left((I + T + M_{\Phi I}) A_\gamma - \gamma_B\right) \gamma_U - k_{U_\gamma} \gamma_B, \tag{17}$$

$$\frac{d\alpha_U}{dt} = \frac{p_{\alpha,T} T}{T + \eta_{\alpha,T}} + \frac{p_{\alpha,M_{\Phi I}} M_{\Phi I}}{M_{\Phi I} + \eta_{\alpha,M_{\Phi I}}} + \frac{p_{\alpha,M} M}{M + \eta_{\alpha,M}} + \frac{p_{\alpha,K} K}{K + \eta_{\alpha,K}} - k_{lin_\alpha} \alpha_U \tag{18}$$

$$- k_{B_\alpha} \left((I + T + M_{\Phi I} + K + M) A_\alpha - \alpha_B\right) \alpha_U + k_{U_\alpha} \alpha_B,$$

$$\frac{d\alpha_B}{dt} = -k_{int_\alpha} \alpha_B + k_{B_\alpha} \left((I + T + M_{\Phi I} + K + M) A_\alpha - \alpha_B\right) \alpha_U - k_{U_\alpha} \alpha_B, \tag{19}$$

$$\frac{dG_U}{dt} = \frac{p_{G,M_{\Phi I}} M_{\Phi I}}{M_{\Phi I} + \eta_{G,M_{\Phi I}}} + \frac{p_{G,M} M}{M + \eta_{G,M}} - k_{lin_G} G_U - k_{B_G} (M A_G - G_B) G_U + k_{U_G} G_B, \tag{20}$$

$$\frac{dG_B}{dt} = -k_{int_G} G_B + k_{B_G} (M A_G - G_B) G_U - k_{U_G} G_B, \tag{21}$$

$$\frac{dC_U}{dt} = \frac{p_{C,M} M}{M + \eta_{C,M}} - k_{lin_C} C_U - k_{B_C} (N A_C - C_B) (C_U)^W + k_{U_C} C_B, \tag{22}$$

$$\frac{dC_B}{dt} = -k_{int_C} C_B + k_{B_C} (N A_C - C_B) (C_U)^W - k_{U_C} C_B. \tag{23}$$

Table 1: List of model variables and their description in Eqs. (1)–(23)

| Variable | Description | Unit |
|---|---|---|
| $V$ | Viral load | cop/ml |
| $S$ | Susceptible cells | $10^9$ cells/ml |
| $I$ | Infected cells | $10^9$ cells/ml |
| $R$ | Resistant cells | $10^9$ cells/ml |
| $D$ | Dead cells | $10^9$ cells/ml |
| $M_{\Phi I}$ | Inflammatory macrophages | $10^9$ cells/ml |
| $M_{\Phi R}$ | Alveolar (or resident) macrophages | $10^9$ cells/ml |
| $K$ | NK cells | $10^9$ cells/ml |
| $T$ | $CD8^+$ T cells | $10^9$ cells/ml |
| $M$ | Monocytes | $10^9$ cells/ml |
| $N$ | Neutrophils | $10^9$ cells/ml |
| $K_R$ | Bone marrow reservoir NK cells | $10^9$ cells/ml |
| $M_R$ | Bone marrow reservoir monocytes | $10^9$ cells/ml |
| $N_R$ | Bone marrow reservoir neutrophils | $10^9$ cells/ml |
| $L_{U,B}$ | Unbound ($U$) or bound ($B$) IL-6 | pg/ml |
| $F_{U,B}$ | Unbound or bound type I IFN | pg/ml |
| $\gamma_{U,B}$ | Unbound or bound IFN-$\gamma$ | pg/ml |
| $\alpha_{U,B}$ | Unbound or bound TNF-$\alpha$ | pg/ml |
| $G_{U,B}$ | Unbound or bound GM-CSF | pg/ml |
| $C_{U,B}$ | Unbound or bound G-CSF | pg/ml |
| $C_{BF}$ | Neutrophil G-CSF receptor bound fraction | unitless |

Table 2: Description of selected biological pathways in Eqs. (1)–(23)

| Pathway | Description |
|---|---|
| $a$ | Cellular lysis |
| $b$ | Destruction of susceptible cells by neutrophils |
| $c$ | Type I IFN and IFN-$\gamma$ inhibition of viral replication |
| $d$ | IL-6 inhibition of infected cells killed by NK cells |
| $e$ | Macrophage exhaustion for dead cells |
| $f$ | Macrophage apoptosis induced by IFN-$\gamma$ and TNF-$\alpha$ |
| $g$ | Monocyte to macrophage differentiation inhibited by TNF-$\alpha$ |
| $h$ | IL-6-induced monocyte differentiation inhibited by TNF-$\alpha$ |
| $i$ | T cell proliferation by type I IFN |
| $j$ | T cell clearance by NK cells, which is inhibited by IL-6 |
| $k$ | Pharmacokinetics and pharmacodynamics of GM-CSF on monocytes |
| $l$ | Production of IL-6 stimulated by infected cells, macrophages, and monocytes |
| $m$ | IL-6 renal clearance |
| $n$ | IL-6 binding |
| $o$ | IL-6 unbinding |
| $p$ | IL-6 internalization |

Table 3: Viral kinetic parameters

| Symbol | Description | Unit | Value | Reference |
|---|---|---|---|---|
| $p$ | Lytic viral production rate | 1/day × $\log_{10}$(cop/ml)/$10^9$ cells | 591 | [1] |
| $\beta$ | Transmission rate (or virus infection rate) | 1/day × 1/$\log_{10}$(cop/ml)/$10^9$ cells | 0.29 | [1] |
| $S_{max}$ | Epithelial cells carrying capacity | $10^9$ cells | $S_0 = 0.16$ | [2] |
| $M_{\Phi_{max}}$ | Macrophage carrying capacity | $10^9$ cells | $M_{\Phi R,0} = 2.7\times10^{-5}$ | [3] |
| $\lambda_S$ | Proliferation of epithelial cells | 1/day | 0.74 | [4] |
| $\lambda_{M_\Phi}$ | Production of alveolar macrophages | $\log_{10}$(cop/ml)/day | 5943 | [5] |
| $\tau_I$ | Eclipse time | day | 0.17 | [6] |
| $d_I$ | Death rate of infected cells | 1/day | 0.1 | [3] |
| $\tau_T$ | Delay in $CD8^+$ T cell arrival | day | 4.5 | [7] |

Table 4: Cell production, recruitment, and activation rates (H refers to homeostasis calculation)

| Symbol | Description | Unit | Value | Reference |
|---|---|---|---|---|
| $p_{M_{\Phi I},G}$ | Monocyte to macrophage differentiation by GM-CSF | 1/day | 1.68 | [8] |
| $p_{M_{\Phi I},L}$ | Monocyte to macrophage differentiation by IL-6 | 1/day | 0.78 | [9, 10] |
| $a_{I,M_\Phi}$ | Activation of macrophages by infected and dead cells | ml/($10^9$ cells) × (1/day) | $1.1 \times 10^3$ | [11, 12] |
| $p_{M,I}$ | Monocyte recruitment rate by infected cells | 1/day | 0.22 | [13] |
| $p_{T,F}$ | $CD8^+$ T cell production rate by IFN | 1/day | 4 | [14] |
| $p_{T,\gamma}$ | $CD8^+$ T cell production rate by IFN-$\gamma$ | 1/day | 6.56 | [15] |
| $p_{N,L}$ | Neutrophil recruitment rate by IL-6 | 1/day | 0.21 | Homeostasis (H) |
| $p_{T,L}$ | $CD8^+$ T cell recruitment rate by IL-6 | 1/day | 4 | [16] |
| $p_{T,I}$ | $CD8^+$ T cell proliferation rate | 1/day | 0.016 | [17] |
| $p_K$ | NK proliferation rate | 1/day | 0.0365 | [18] |
| $p_{K,\alpha}$ | NK production rate by TNF-$\alpha$ | 1/day | 0.2148 | [18] |
| $M^*_{prod}$ | Homeostasis reservoir release rate | 1/day | 0.13 | H |
| $\psi_{M_{max}}$ | Maximal reservoir release rate | 1/day | 11.55 | [19] |
| $N^*_{prod}$ | Homeostasis reservoir release rate | 1/day | 0.21 | H |
| $\psi_{N_{max}}$ | Maximal reservoir release rate | 1/day | 4.13 | [20] |
| $K^*_{prod}$ | Homeostasis reservoir release rate | 1/day | 0.1 | H |
| $C^*_{BF}$ | Homeostasis neutrophil receptor bound fraction | – | $1.6 \times 10^{-5}$ | [20] |

### 2. Parameters taken from literature

#### *2.1. NK cell proliferation rate*

Natural Killer (NK) cells are proliferated and activated by various cytokines that are not explicitly represented in our model. For example, IL-2 is a primary cytokine responsible for the proliferation and activation of NK cells. To make up for IL-2's effect, incorporating a proliferation rate into Eq. (8) in our model is essential. The proliferation rate of NK cells in both young and elderly adults was measured using Ki67 assay [43]. The results indicated no significant difference between the two age groups, so the average NK proliferation rate of $p_K = 0.0365$/day.

#### *2.2. NK cell death rate*

Like normal cells, NK cells undergo cell death via apoptosis. They undergo apoptosis to regulate immune responses, ensure immune system balance, and prevent prolonged or excessive activity. Hence, a death rate for NK is necessary in Eq. (8). The apotosis rate of NK cells in both young and elderly adults was measured using TUNEL assay [43]. The results similarly indicated no significant difference between the two age groups, so the average NK apoptosis rate of $d_K = 0.156$/day is used.

Table 5: Cell-related half-effect ($\epsilon$), IC50 ($IC_{50}$), and Hill coefficient ($h$) parameters

| Symbol | Description | Unit | Value | Reference |
|---|---|---|---|---|
| $\epsilon_{F,I}$ | Type I IFN inhibition of viral production | pg/ml | $2 \times 10^{-4}$ | [21] |
| $\epsilon_{\gamma,I}$ | IFN-$\gamma$ inhibition of viral production | pg/ml | $3.147 \times 10^{-4}$ | [22] |
| $\epsilon_{L,M_\Phi}$ | IL-6 monocytes to macrophages | pg/ml | 0.001 | [10] |
| $\epsilon_{\alpha,M_{\Phi I}}$ | TNF-$\alpha$ inhibition of macrophage differentiation | pg/ml | 0.001093 | [10] |
| $\epsilon_{G,M_{\Phi I}}$ | GM-CSF monocyte to macrophages | pg/ml | 0.027 | [23] |
| $\epsilon_{G,M}$ | GM-CSF recruitment of monocytes | pg/ml | 57.2 | [24] |
| $\epsilon_{F,T}$ | Type I IFN production of $CD8^+$ T cells | pg/ml | 0.004 | [25] |
| $\epsilon_{\gamma,T}$ | IFN-$\gamma$ production of $CD8^+$ T cells | pg/ml | 0.004 | [25] |
| $\epsilon_{C,N}$ | G-CSF recruitment of neutrophils | unitless | $1.89 \times 10^{-4}$ | [20] |
| $\epsilon_{L,N}$ | IL-6 recruitment of neutrophils | pg/ml | 57.2 | [24] |
| $\epsilon_{I,M}$ | Infected cell monocyte recruitment | $10^9$ cells/ml | 0.11 | [3] |
| $\epsilon_{L,T}$ | IL-6 production of $CD8^+$ T cells | pg/ml | $1.5 \times 10^{-5}$ | [26] |
| $\epsilon_{V,M_\Phi}$ | Viral load for macrophage replenishing | $\log_{10}$(cop/ml) | 0.905 | [27] |
| $\epsilon_{T,I}$ | Antigen driven proliferation | $10^9$ cells/ml | $10^{-6}$ | [17] |
| $\epsilon_{L,K}$ | IL-6 inhibition of NK cell cytotoxicity | pg/ml | $2.102 \times 10^{-5}$ | [28] |
| $\epsilon_{K,I}$ | NK cells lyse infected cells | $10^9$ cells/ml | $2.345\times 10^{-3}$ | [29] |
| $\epsilon_{K,T}$ | NK cells lyse T cells | $10^9$ cells/ml | $3.646\times 10^{-3}$ | [30] |
| $\epsilon_{\alpha,K}$ | TNF-$\alpha$ recruitment of NK cells | pg/ml | 0.06192 | [18] |
| $\epsilon_{\gamma,\alpha}$ | TNF-$\alpha$ and IFN-$\gamma$ synergestic induced damage to inflammatory macrophages | $(pg/ml)^2$ | $1.994 \times 10^{-5}$ | [31] |
| $h_M$ | GM-CSF monocyte recruitment | – | 1.67 | [24] |
| $h_{M_{\Phi I},\alpha}$ | TNF-$\alpha$ inhibition of inflammatory macrophage differentiation | – | 0.4 | [10] |
| $h_{M,M_\Phi}$ | GM-CSF monocyte to macrophages | – | 2.03 | [23] |
| $h_N$ | Neutrophil induced damage | – | 3.02 | [11] |
| $IC_{50,N}$ | Neutrophil induced damage | $10^9$ cells/ml | 0.047 | [11] |

Table 6: Cell/virus-induced death rates

| Symbol | Description | Unit | Value | Reference |
|---|---|---|---|---|
| $\delta_{V,M_\Phi}$ | Rate of viral clearance by macrophages | ml/($10^9$ cells) × 1/day | 1152 | [32] |
| $\delta_{V,N}$ | Rate of viral clearance by neutrophils | ml/($10^9$ cells) × 1/day | 1047 | [3] |
| $\delta_N$ | Rate of neutrophil inflicted damage | 1/day | 1.68 | [32] |
| $\rho$ | Bystander death modulation constant | – | 0.5 | [3] |
| $\delta_{I,M_\Phi}$ | Rate macrophages phagocytose infected cells | ml/($10^9$ cells) × 1/day | 121 | [33] |
| $\delta_{I,T}$ | Rate $CD8^+$ T cells induce apoptosis in infected cells | ml/($10^9$ cells) × 1/day | 119 | [34] |
| $\delta_{I,K}$ | Rate NK cells lyse infected cells | 1/day | 1.037 | [29] |
| $\delta_{T,K}$ | Rate NK cells lyse T cells | 1/day | 0.1593 | [30] |
| $\delta_{M_{\Phi I}}$ | Rate inflammatory macrophages die due to TNF-$\alpha$ and IFN-$\gamma$ | 1/day | 0.1543 | [31] |
| $\delta_{M_{\Phi,D}}$ | Rate macrophages die from phagocytosis | ml/($10^9$ cells) × 1/day | 6.06 | [35, 36] |
| $\delta_{D,M_\Phi}$ | Rate macrophages phagocytose dead cells | ml/($10^9$ cells) × 1/day | 8.03 | [35] |

#### 2.3. TNF-$\alpha$ internalization rate

Similarly to other cytokines, TNF-$\alpha$ occasionally internalizes into a cell after binding to cell receptors [98]. It should be noted that TNF-$\alpha$ can contain two distinct internalization rates due to having multiple receptors. Namely, TNF-$\alpha$ can bind to TNF-$\alpha$ receptor 1 (TNFR1) and TNF-$\alpha$ receptor 2 (TNFR2) [67]. TNFR1 and TNFR2 have internalization rates of 66.528 and 39.744 per day, respectively [70]. To capture both receptors in our model while

Table 7: Cell death and virus decay rates

| Symbol | Description | Unit | Value | Reference |
|---|---|---|---|---|
| $d_V$ | Viral decay rate | 1/day | 0 | [37] |
| $d_D$ | Degradation rate of apoptosed cells | 1/day | 8 | [38] |
| $d_{M_{\Phi R}}$ | Alveolar macrophage death rate | 1/day | 0 | [39] |
| $d_{M_{\Phi I}}$ | Inflammatory macrophage death rate | 1/day | 0.3 | [40] |
| $d_M$ | Monocyte death rate | 1/day | 0.76 | [41] |
| $d_N$ | Neutrophil death rate | 1/day | 1.28 | [20] |
| $d_T$ | $CD8^+$ T cell death rate | 1/day | 0.4 | [42] |
| $d_K$ | NK cell death rate | 1/day | 0.156 | [43] |

Table 8: Cytokine production rates

| Symbol | Description | Unit | Value | Reference |
|---|---|---|---|---|
| $p_{L,I}$ | IL-6 production by infected cells | pg/ml/day | 11.89 | [44] |
| $p_{L,M_\Phi}$ | IL-6 production by activated macrophages | pg/ml/day | 1872 | [45] |
| $p_{L,M}$ | IL-6 production by monocytes | pg/ml/day | 36.28 | [46] |
| $p_{G,M_{\Phi I}}$ | GM-CSF production by inflammatory macrophages | pg/ml/day | 2626 | [47] |
| $p_{G,M}$ | GM-CSF production by monocytes | pg/ml/day | 3070 | [48] |
| $p_{C,M}$ | G-CSF production by monocytes | pg/ml/day | 30.70 | [3] |
| $p_{F,I}$ | Type I IFN production by infected cells | pg/ml/day | 2.82 | [49] |
| $p_{F,M_{\Phi I}}$ | Type I IFN production by inflammatory macrophages | pg/ml/day | 1.3 | [3] |
| $p_{F,M}$ | Type I IFN production by monocytes | pg/ml/day | 3.56 | [50, 51] |
| $p_{\gamma,K}$ | IFN-$\gamma$ production by NK cells | pg/ml/day | 325 | [52] |
| $p_{\gamma,T}$ | IFN-$\gamma$ production by T cells | pg/ml/day | 300 | [53] |
| $p_{\alpha,M_{\Phi I}}$ | TNF-$\alpha$ production by inflammatory macrophages | pg/ml/day | 3824 | [45] |
| $p_{\alpha,M}$ | TNF-$\alpha$ production by monocytes | pg/ml/day | 110 | [54, 55] |
| $p_{\alpha,T}$ | TNF-$\alpha$ production by T cells | pg/ml/day | 450 | [56] |
| $p_{\alpha,K}$ | TNF-$\alpha$ production by NK Cells | pg/ml/day | 1000 | [52, 57] |

incorporating the fact that TNFR1 having higher affinity [67], we take a weighted average of both internalization rates, with TNFR1 having a weight of 0.66 and TNFR2 having a weight 0.33. Through this process we obtain a weighted average $k_{int_\alpha}$ of 57.6 per day.

#### 2.4. *TNF-α unbinding and binding, and renal clearance rates*

Upon binding to a cell receptor, cytokines will eventually unbind, discontinuing the cytokine to cell interaction [78]. TNF-$\alpha$'s unbinding average across both TNFR1 and TNFR2 receptors are found to be 449.46 per day [70]. We thus use $k_{U_\alpha}$ = 449.46/day. The rate in which TNF-$\alpha$ binds to cell receptors is found by calculating the weighted average between both receptors. Using different weights (0.66 and 0.33) for TNFR1 and TNFR2, respectively, our average binding rate is $k_{B_\alpha}$ = 0.096 pg/ml/day [70].

Circulating (unbound) TNF-$\alpha$ is filtered and cleared away from the system at rate $k_{lin_\alpha}$ of 33.27 per day [62].

#### 2.5. *TNF-α molecular weight and receptor counts*

TNF-$\alpha$, in its common form, has been reported by various studies [85, 86] to possess a molecular weight of 17,300 kD (or g/mol). Thus, $MM_\alpha$ = 17,300 g/mol.

In our model, monocytes, inflammatory macrophages, NK cells, T cells, and infected cells are found to have significant numbers of TNF-$\alpha$ receptors (TNFR) [70]. While some studies have varying numbers of TNFR, with an upper and lower bound of receptor counts, we specifically looked at lower bounds as we found that studies with

Table 9: Cytokine production half-effect ($\eta$) parameters

| Symbol | Description | Unit | Value | Reference |
|---|---|---|---|---|
| $\eta_{L,I}$ | IL-6 production by infected cells | $10^9$ cells/ml | 0.7 | [44] |
| $\eta_{L,M}$ | IL-6 by monocytes | $10^9$ cells/ml | 0.0045 | [46] |
| $\eta_{L,M_{\Phi I}}$ | IL-6 by inflammatory macrophages | $10^9$ cells/ml | $1.82 \times 10^{-4}$ | [45] |
| $\eta_{G,M_{\Phi I}}$ | GM-CSF by macrophages | $10^9$ cells/ml | $1.82 \times 10^{-4}$ | [3] |
| $\eta_{G,M}$ | GM-CSF by monocytes | $10^9$ cells/ml | 0.15 | H |
| $\eta_{C,M}$ | G-CSF by monocytes | $10^9$ cells/ml | $8 \times 10^{-4}$ | H |
| $\eta_{F,I}$ | Type I IFN by infected cells | $10^9$ cells/ml | 0.011 | [49] |
| $\eta_{F,M_{\Phi I}}$ | Type I IFN by inflammatory macrophages | $10^9$ cells/ml | $2 \times 10^{-4}$ | H |
| $\eta_{F,M}$ | Type I IFN by monocytes | $10^9$ cells/ml | 0.54 | [50, 51] |
| $\eta_{\gamma,K}$ | IFN-$\gamma$ by NK cells | $10^9$ cells/ml | $2.99\times10^{-4}$ | [52] |
| $\eta_{\gamma,T}$ | IFN-$\gamma$ by T cells | $10^9$ cells/ml | $8.37\times10^{-5}$ | [41] |
| $\eta_{\alpha,M_{\Phi I}}$ | TNF-$\alpha$ by inflammatory macrophages | $10^9$cells/ml | $2.22\times10^{-4}$ | [58] |
| $\eta_{\alpha,M}$ | TNF-$\alpha$ by monocytes | $10^9$cells/ml | 0.3851 | [54, 55] |
| $\eta_{\alpha,T}$ | TNF-$\alpha$ by T cells | $10^9$cells/ml | $9.706\times 10^{-5}$ | [56] |
| $\eta_{\alpha,K}$ | TNF-$\alpha$ by NK cells | $10^9$cells/ml | $4.29 \times 10^{-4}$ | [52, 57] |

Table 10: Cytokine (renal) clearance and internalization rates

| Symbol | Description | Unit | Value | Reference |
|---|---|---|---|---|
| $k_{lin_L}$ | Rate of IL-6 renal clearance | 1/day | 16.6 | [59] |
| $k_{lin_G}$ | Rate of GM-CSF renal clearance | 1/day | 11.7 | [60] |
| $k_{lin_C}$ | Rate of G-CSF renal clearance | 1/day | 0.16 | [20] |
| $k_{lin_F}$ | Rate of type I IFN renal clearance | 1/day | 16.63 | [61] |
| $k_{lin_\gamma}$ | Rate of IFN-$\gamma$ renal clearance | 1/day | 28.5 | [61] |
| $k_{lin_\alpha}$ | Rate of TNF-$\alpha$ renal clearance | 1/day | 33.27 | [62] |
| $k_{int_L}$ | Internalization rate of IL-6 | 1/day | 61.8 | [63] |
| $k_{int_G}$ | Internalization rate of GM-CSF | 1/day | 73.4 | [64] |
| $k_{int_C}$ | Internalization rate of G-CSF | 1/day | 462 | [20] |
| $k_{int_F}$ | Internalization rate of type I IFN | 1/day | 17 | [65] |
| $k_{int_\gamma}$ | Internalization rate of IFN-$\gamma$ | 1/day | 17 | [3] |
| $k_{int_\alpha}$ | Internalization rate of TNF-$\alpha$ | 1/day | 57.6 | [66, 67] |

higher levels of TNFR often contained stimulants that likely altered surface receptor expressions [78]. Concluding our search for TNF-$\alpha$ receptor counts, we find that approximately, monocytes contain 230 TNFR, CD8$^+$ T cells contain 300 TNFR, infected cells contain 714 TNFR, and macrophages contain 1500 TNFR [70]. Since it is difficult to find NK cell receptor counts, we decide to set NK TNFR expression similar to monocyte TNFR expressions as both are from the same lineage of cells [78].

#### 2.6. TNF-$\alpha$ initial unbound and bound levels

We obtained the initial levels of unbounded TNF-$\alpha$ by finding the average circulating serum levels of TNF-$\alpha$ in healthy adults across diverse age groups [97]. Studies suggest unbounded levels to be around 0.7 pg/ml, with a statistical significance of $p < 0.025$. Regarding the bounded TNF-$\alpha$ parameter, we calculated it using the following equations:

$$A_\alpha = \frac{MM_\alpha \times 10^{12}}{6.02214 \times 10^{-23}}(K_{\alpha,M_{\Phi I}} + K_{\alpha,T} + K_{\alpha,I} + K_{\alpha,M} + K_{\alpha,K}) \times 10^9 \times \frac{1}{5000},$$

Table 11: Cytokine binding/unbinding rates and stoichiometric constants

| Symbol | Description | Unit | Value | Reference |
|---|---|---|---|---|
| $k_{B_L}$ | IL-6 binding rate | ml/pg/day | 0.0018 | [68] |
| $k_{B_G}$ | GM-CSF binding rate | ml/pg/day | 0.0021 | [64] |
| $k_{B_C}$ | G-CSF binding rate | ml/ng/day | 2.24 | [20] |
| $k_{B_F}$ | IFN binding rate | ml/pg/day | 0.011 | [65] |
| $k_{B_\gamma}$ | IFN-$\gamma$ binding rate | ml/pg/day | 0.0382 | [69] |
| $k_{B_\alpha}$ | TNF-$\alpha$ binding rate | ml/pg/day | 0.096 | [70] |
| $k_{U_L}$ | IL-6 unbinding rate | 1/day | 22.3 | [68] |
| $k_{U_G}$ | GM-CSF unbinding rate | 1/day | 522 | [64] |
| $k_{U_C}$ | G-CSF unbinding rate | 1/day | 184 | [20] |
| $k_{U_F}$ | IFN unbinding rate | 1/day | 6.07 | [65] |
| $k_{U_\gamma}$ | IFN-$\gamma$ unbinding rate | 1/day | 432 | [69] |
| $k_{U_\alpha}$ | TNF-$\alpha$ unbinding rate | 1/day | 449.46 | [70] |
| W | Stoichiometric constant (G-CSF) | – | 1.4608 | [20] |
| $\hat{p}$ | Stoichiometry relating constant (IL-6, GM-CSF, IFN, IFN-$\gamma$, TNF-$\alpha$) | – | 1 | [3] |
| $\hat{p}$ | Stoichiometry relating constant (G-CSF) | – | 2 | [20] |

Table 12: Number of cellular receptors and cytokine molecular weights

| Symbol | Description | Unit | Value | Reference |
|---|---|---|---|---|
| $K_{L,N}$ | No. IL-6 receptors on neutrophils | sites/cell | 720 | [71] |
| $K_{L,T}$ | No. IL-6 receptors on T cells | sites/cell | 300 | [72] |
| $K_{L,M}$ | No. IL-6 receptors on monocytes | sites/cell | 509 | [73] |
| $K_{G,M}$ | No. GM-CSF receptors on monocytes | sites/cell | 1058 | [74] |
| $K_{C,N}$ | No. G-CSF receptors on neutrophils | sites/cell | 600 | [75] |
| $K_{F,T}$ | No. of type I IFN receptors on T cells | sites/cell | 1000 | [76] |
| $K_{F,I}$ | No. of type I IFN receptors on infected cells | sites/cell | 1300 | [77] |
| $K_{\gamma,T}$ | No. of IFN-$\gamma$ receptors on T cells | sites/cell | 500 | [69] |
| $K_{\gamma,I}$ | No. of IFN-$\gamma$ receptors on infected cells | sites/cell | 1800 | [69] |
| $K_{\gamma,M_{\Phi I}}$ | No. of IFN-$\gamma$ receptors on inflammatory macrophages | sites/cell | 760 | [76] |
| $K_{\alpha,M}$ | No. of TNF-$\alpha$ receptors on monocytes | sites/cell | 230 | [78] |
| $K_{\alpha,T}$ | No. of TNF-$\alpha$ receptors on T cells | sites/cell | 300 | [70] |
| $K_{\alpha,M_{\Phi I}}$ | No. of TNF-$\alpha$ receptors on inflammatory macrophages | sites/cell | 1500 | [70] |
| $K_{\alpha,I}$ | No. of TNF-$\alpha$ receptors on infected cells | sites/cell | 714 | [79] |
| $K_{\alpha,K}$ | No. of TNF-$\alpha$ receptors on NK cells | sites/cell | 230 | [78] |
| $MM_L$ | Molecular weight of IL-6 | g/mol | 21000 | [80, 81, 82] |
| $MM_G$ | Molecular weight of GM-CSF | g/mol | 14000 | [83] |
| $MM_C$ | Molecular weight of G-CSF | g/mol | 19600 | [20] |
| $MM_F$ | Molecular weight of IFN-$\beta$ | g/mol | 19000 | [84] |
| $MM_\gamma$ | Molecular weight of IFN-$\gamma$ | g/mol | 16500 | [61] |
| $MM_\alpha$ | Molecular weight of TNF-$\alpha$ | g/mol | 17300 | [85, 86] |

$$\alpha_{B_0} = \frac{k_B \times A_\alpha \times \alpha_{U_0} \times (M_{\Phi I} + M + K)}{k_B \times \alpha_{U_0} + k_{int} + k_U}.$$

Using Eq. (46) (calculated from homeostasis), we found initial bounded TNF-$\alpha$ levels $\alpha_{B,0}$ to be approximately $1.3 \times 10^{-6}$ pg/ml.

Table 13: Initial conditions

| Symbol | Description | Unit | Value | Reference |
|---|---|---|---|---|
| $V_0$ | Initial viral load | $\log_{10}$(cop/ml) | 4.5 | [1] |
| $S_0$ | Initial susceptible cells | $10^9$ cells/ml | 0.16 | [2], [87] |
| $I_0$ | Initial infected cells | $10^9$ cells/ml | 0 | |
| $R_0$ | Initial resistant cells | $10^9$ cells/ml | 0 | |
| $M_{\Phi R,0}$ | Initial resident macrophages | $10^9$ cells/ml | $2.7 \times 10^{-5}$ | [2] |
| $M_{\Phi I,0}$ | Initial inflammatory macrophages | $10^9$ cells/ml | $1.5 \times 10^{-6}$ | H |
| $M_0$ | Initial monocytes | $10^9$ cells/ml | 0.0004 | [88] |
| $M_R$ | Initial reservoir monocytes | $10^9$ cells/ml | 0.0023 | [3] |
| $N_0$ | Initial neutrophils | $10^9$ cells/ml | 0.0053 | [20] |
| $N_R$ | Initial reservoir neutrophils | $10^9$ cells/ml | 0.0316 | [20] |
| $T_0$ | Initial CD8$^+$ T cells | $10^9$ cells/ml | $1.1 \times 10^{-4}$ | [89, 90] |
| $K_0$ | Initial NK cells | $10^9$ cells/ml | $7.15 \times 10^{-5}$ | [91] |
| $K_R$ | Initial reservoir NK cells | $10^9$ cells/ml | $1.1 \times 10^{-4}$ | [92] |
| $L_{U,0}$ | Initial unbound IL-6 | pg/ml | 1.1 | [93] |
| $L_{B,0}$ | Initial bound IL-6 | pg/ml | $1.4 \times 10^{-6}$ | H |
| $G_{U,0}$ | Initial unbound GM-CSF | pg/ml | 2.43 | [94] |
| $G_{B,0}$ | Initial bound GM-CSF | pg/ml | $1.6 \times 10^{-8}$ | H |
| $C_{U,0}$ | Initial unbound G-CSF | ng/ml | 0.025 | [20] |
| $C_{B,0}$ | Initial bound G-CSF | ng/ml | $6.5 \times 10^{-10}$ | [20] |
| $F_{U,0}$ | Initial unbound type I IFN | pg/ml | 0.015 | [95] |
| $F_{B,0}$ | Initial bound type I IFN | pg/ml | $1.1 \times 10^{-8}$ | H |
| $\gamma_{U,0}$ | Initial unbound IFN-$\gamma$ | pg/ml | 0.91 | [96] |
| $\gamma_{B,0}$ | Initial bound IFN-$\gamma$ | pg/ml | $1.432 \times 10^{-7}$ | H |
| $\alpha_{U,0}$ | Initial unbound TNF-$\alpha$ | pg/ml | 0.7 | [97] |
| $\alpha_{B,0}$ | Initial bound TNF-$\alpha$ | pg/ml | $1.3 \times 10^{-6}$ | H |

Table 14: Parameter modifications for severe trend line

| Symbol | Description | Unit | Mild Case | Severe Case | Reference |
|---|---|---|---|---|---|
| $p_{F,I}$ | Production of type I IFN by Infected Cells | pg/ml/day | 2.82 | 0.002 | [49] |
| $p_{M,I}$ | Monocyte recruitment rate by infected cells | 1/day | 0.22 | 1.2 | [13] |
| $\eta_{F,M\Phi I}$ | Cytokine production half-effect coefficient of type I IFN by inflammatory macrophages | $10^9$ cells/ml | $1.3 \times 10^{-6}$ | $2 \times 10^{-4}$ | H |
| $\epsilon_{L,K}$ | Half effect constant for IL-6 inhibition of NK cell cytotoxicity | pg/ml | $2.102 \times 10^{-5}$ | $2.102 \times 10^{-6}$ | [28] |
| $p_{\gamma,T}$ | Production rate of IFN-$\gamma$ by T cells | pg/ml/day | 300 | 30 | [53] |

#### *2.7. IFN-$\gamma$ inhibition of viral production*

Both type I IFN and IFN-$\gamma$ inhibit viral production through the release of Interferon Stimulated Genes (ISGs) [99]. The study in [22] attempted to measure how well IFN-$\gamma$ could inhibit viral replication as well as type 1 IFNs. Using viral plaques as the measure of viral inhibition, this study showed that 100 U/ml of IFN-$\gamma$ was enough to cut viral plaque numbers in half, representing a half effect concentration. In order to convert this number to pg/ml, two steps were taken. First, we divide the half effect concentration by a specific activity derived from the results in [100]. This provides us with $\frac{100}{5\times10^7} = 2000$ pg/ml.

Since 2000 pg/ml is the number of free circulating IFN-$\gamma$ that produces this effect, we must convert this number to the amount of bound IFN-$\gamma$. We do this by multiplying the unbounded half effect by the ratio between initial bound and unbound IFN-$\gamma$. That is, $2000 \times \frac{1.432\times10^{-7}}{0.91} = 3.147 \times 10^{-4}$ pg/ml.

#### 2.8. *Internalization rate of IFN-$\gamma$*

Given limited information on the pharamcodynamics of IFN-$\gamma$, we choose to use the internalization rate of IFN type I, reported by [3] to be 17/day.

#### 2.9. *IFN-$\gamma$ unbinding and binding, and renal clearance rates*

The half life of IFN-$\gamma$ was reported by [61] to be between 25-35 minutes. Picking the upper bound gives us a half life equivalent to 0.0243 days. We then turned the half life into the clearance rate by re-arranging equations from [101]. This gives us $k_{lin_\gamma} = \frac{ln(2)}{0.0243} = 28.5$/day. Sadir et al. [69] reported an unbinding rate $k_{\text{off}} = 5 \times 10^{-3}$/s for IFN-$\gamma$, which converts to $k_{U_\gamma} = 0.005 \times 86{,}400 = 432$/day.

The reported binding rate is $k_{\text{on}} = 7.3 \times 10^6$ 1/(M × s) (or L/(mol × s)) [69]. To express this in ml/pg/day, we divide by the molecular weight $M_\gamma$ in ml/pg/day and convert units: $k_{\text{on}} = (7.3 \times 10^6 \times 1000 \times 86{,}400)/M_\gamma$. We then derive the molar mass from the molecular weight, taking $M_\gamma = 16.5$ kDa $= 1.65 \times 10^{16}$ pg mol$^{-1}$ [102], giving $k_{B_\gamma} \approx 0.0382$ ml/pg/day.

Lastly, the terminal half-life of human IFN-$\gamma$ is $t_{1/2} = 25$–$35$ min [61]; using the upper bound (35 min = 0.0243 day), the first-order clearance rate is $k_{lin_\gamma} = \ln(2)/t_{1/2} \approx 28.5$/day.

#### 2.10. *IFN-$\gamma$ receptor counts*

The amount of IFN-$\gamma$ receptors on T cells and infected cells are found from [69]. The receptor counts are $K_{\gamma,I} = 1800$ receptors/cell and $K_{\gamma,T} = 500$ receptors/cells. The receptor counts for macrophages were found to be $K_{\gamma,M_{\Phi I}} = 760$ receptors/cell from [76].

### 3. Parameters estimated from data fitting

Figs. 1–3 present the parameter estimation results obtained through data fitting. Detailed explanations are provided in each subsection below.

#### 3.1. *Effect of NK cells on infected cells*

At higher relative densities of target cells, it is more likely that NK cells encounter target cells that can trigger degranulation. This means that the number of infected cells affect the NK cell's cytotoxicity. NK cell's targeting on infected cells is therefore modeled by:

$$\frac{dI}{dt} = -\frac{\delta_{I,K} I K}{K + \epsilon_{K,I}}, \tag{24}$$

where $\delta_{I,K}$ is the infected cells death rate by NK cells measured in 1/day. The study in [29] measured NK cell lysis dependent on NK cell and infected cells. We fit the data to Eq. (24), resulting in $\delta_{I,K} = 1.037$ 1/day and $\epsilon_{K,I} = 2.345 \times 10^{-3}$ (in $10^9$ cells/ml) (shown in Fig. 1A).

#### 3.2. *Effect of NK cells on T cells*

The study in [103] observed an increase in T cells differentiation when NK is depleted in vivo setting. The dynamic is modeled by:

$$\frac{dT}{dt} = -\frac{\delta_{T,K} T K}{K + \epsilon_{K,T}}, \tag{25}$$

where $\delta_{T,K}$ is T-cell death rate by NK cells measured in 1/day. Cytotoxicity assays are conducted utilizing IL-2–activated NK cells as effectors and T cells as target cells [30]. Data fitting involving both NK2D and DNAM1 receptors to the model equation (25) yielded $\delta_{T,K} = 0.1593$ 1/day (Fig. 1B).

#### 3.3. Effect of TNF-$\alpha$ on NK proliferation

TNF-$\alpha$ promotes aerobic glycolysis, mediating NK cell proliferation. Proliferation of NK cells by TNF-$\alpha$ is given by:

$$\frac{dK}{dt} = \frac{p_{K,\alpha}\alpha_B K}{\alpha_B + \epsilon_{\alpha,K}}, \tag{26}$$

where $p_{K,\alpha}$ is the NK production rate by TNF-$\alpha$ measured in 1/day. Khan et al. [18] observed that TNF-$\alpha$ promotes activation and proliferation of NK cells. The percentage of CD25+ cells were measured using stimulation with different doses of TNF-$\alpha$ ex vivo for 3 days. CD25+ is a marker for activated NK cells. Data fitting to the modeled equation yields $p_{K,\alpha} = 0.2148$ 1/day. The fitting is shown in Fig. 1C.

#### 3.4. NK cytotoxicity inhibition by IL-6

IL-6 does not reduce the number of NK cells but decreases their cytotoxic activity. A study in [104] observed no significant difference in the number of NK cells activated with or without IL-6. However, in the presence of IL-6, there was a significant decrease in NK cytotoxicity. This suggests an inhibitory dynamic, where IL-6 reduces the cytotoxicity of NK cells toward infected cells and T-cells. Consequently, the IL-6 term is incorporated into the death term of these cells as mediated by NK cells, modifying Eqs. (24)–(25):

$$\frac{dI}{dt} = -\left(\frac{\delta_{I,K} IK}{K + \epsilon_{K,I}}\right)\left(\frac{\epsilon_{L,K}}{\epsilon_{L,K} + L_B}\right), \tag{27}$$

$$\frac{dT}{dt} = -\left(\frac{\delta_{T,K} TK}{K + \epsilon_{K,T}}\right)\left(\frac{\epsilon_{L,K}}{\epsilon_{L,K} + L_B}\right), \tag{28}$$

where $\epsilon_{L,K}$ is the half coefficient for the inhibition of NK cells by IL-6 measured in pg/ml.

In vivo observation supports that IL-6 reduces the cytolytic activity of NK cells, accompanied by the down-regulation of granzyme B and perforin. A negative correlation between the concentration of IL-6 and the cytolytic activity of NK cells in PF cells was observed and measured [104]. Using the parameters estimated previously for $\delta_{I,K}$ and $\epsilon_{I,K}$, the data of this correlation is used to fit to the equation for infected cells and yielded $\epsilon_{L,K} = 2.102 \times 10^{-5}$ pg/ml.

We conducted additional validation using data [28] by graphing the effect of NK on infected cells with different doses of IL-6 (Fig. 1D). Moreover, infected cells killed by NK cells were predicted with different IL-6 concentrations, as shown in Fig. 1E, to confirm the estimated parameter values.

#### 3.5. TNF-$\alpha$ production by CD8$^+$ T cells

In order to model the production of TNF-$\alpha$ by CD8$^+$ T cells, we used the following fit:

$$\frac{d\alpha_B}{dt} = \frac{p_{\alpha,T} T}{T + \eta_{\alpha,T}}. \tag{29}$$

The production rate was found from the temporal production of TNF-$\alpha$ by activated CD8$^+$ T cells. This yielded a value of 900 pg/ml/day/2 = $p_{\alpha,T}$ = 450 pg/ml/day. The half-effect of production $\eta_{\alpha,T} = 9.706 \times 10^{-5}$ (in $10^9$ cells/ml) was found using a combination of methods. First, we used data from [56], which states the mean fluorescence of TNF-$\alpha$ in a CD8$^+$ T cell culture. Given that mean fluorescence is in RFU (relative fluorescence units), we fit the fluorescence values to a standard curve for TNF-$\alpha$. These values are then fitted using the term below:

$$\frac{d\alpha_B}{dt} = time \times \frac{p_{\alpha,T} T}{T + \eta_{\alpha,T}}. \tag{30}$$

The standard curve and fits are shown in Fig. 2A.

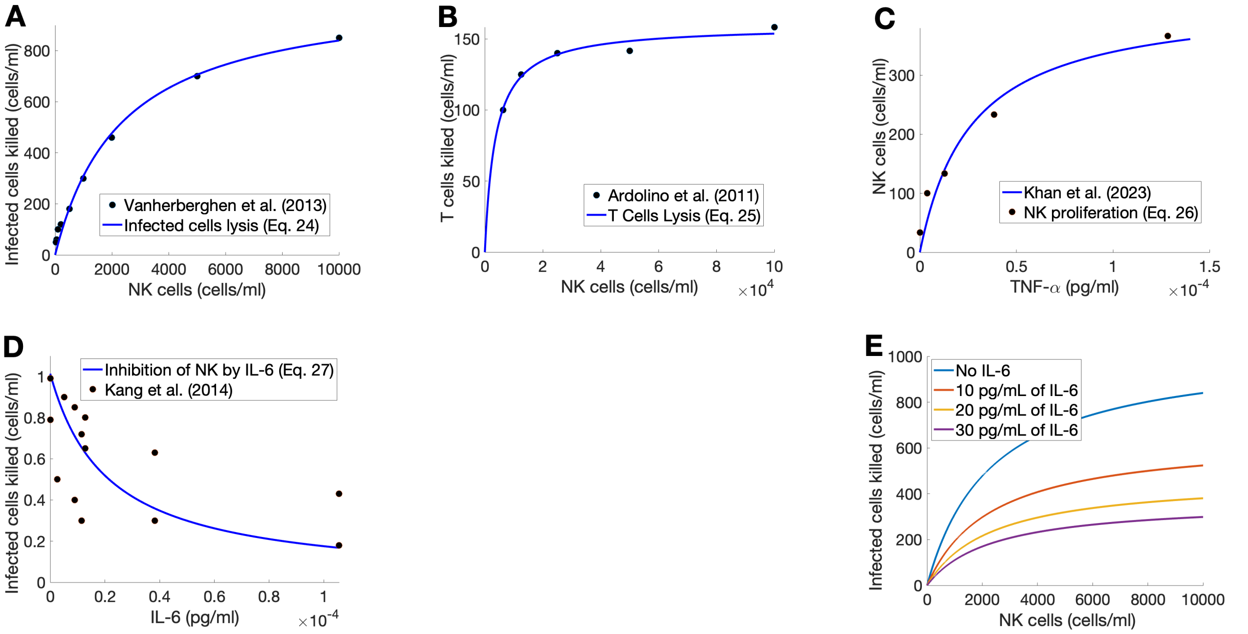


Figure 1: **Dynamics of NK cells on other immune cells and cytokines. A)** Death rate of infected cells by NK cells (Eq. (24)), **B)** Death rate of T cells by NK cells (Eq. (25)), **C)** Proliferation of NK cells by TNF-$\alpha$ (Eq. (26)), **D)** Inhibition of NK cytotoxicity by IL-6 was fitted to estimate the half-coefficient of the inhibition term (Eq. (27)), and **E)** Infected cells killed by NK cells were predicted with different IL-6 concentrations for further confirmation.

#### *3.6. TNF-$\alpha$ production by NK cells*

Similarly to Monocytes, TNF-$\alpha$ secretion by NK cells is also found through data fitting. In the instance of NK cells, data found in [52, 57] is used for data fit to temporal TNF-$\alpha$ production by NK cells. The following fit is shown below:

$$\frac{d\alpha_B}{dt} = \frac{p_{\alpha,K} K}{K + \eta_{\alpha,K}}. \tag{31}$$

Through our findings, we discover that inactive NK cells secrete very little TNF-$\alpha$ levels [52, 57]. To gather accurate pathogenic behaviors in COVID-19 cases, we utilize studies where NK cells were activated by commonly found stimulants in the body [52]. This leads us to a maximum rate $p_{\alpha,K}$ of approximately 1000 pg/ml/day and a half effect $\eta_{\alpha,K}$ of 4.29 $\times 10^{-4}$ (in $10^9$ cells/ml), as demonstrated in Fig. 2B.

#### *3.7. TNF-$\alpha$ inhibition of monocyte to macrophage differentiation*

TNF-$\alpha$ is known to inhibit IL-6 induced monocyte to macrophage differentiation [105]. To capture this behavior, the modified equation from [3] is used, as follows:

$$\frac{dM_{\Phi I}}{dt} = \left(\frac{p_{M_{\Phi I},L} L_B M}{L_B + \epsilon_{L,M_{\Phi I}}}\right)\left(\frac{\epsilon_{\alpha,M_{\Phi I}}^{h_{M_{\Phi I},\alpha}}}{\epsilon_{\alpha,M_{\Phi I}}^{h_{M_{\Phi I},\alpha}} + \alpha_B^{h_{M_{\Phi I},\alpha}}}\right). \tag{32}$$

The Hill coefficient in Eq. 32 above was fit to data found in [105]. Testing with a Hill coefficient proved to give a better fit. However, when fitting our equation, we alterd the original parameters from [3], that is, $p_{M_{\Phi I},L} = 0.78$ 1/day, $\epsilon_{L,M_{\Phi I}} = 0.001$ pg/ml, $\epsilon_{\alpha,M_{\Phi I}} = 0.001093$ pg/ml, and $h_{M_{\Phi I},\alpha} = 0.4$. Thus, to validate the behavior of our parameters, we fit the equation against data sets used in [3]. By doing so, we obtained a great fit when no TNF-$\alpha$ was present ($\alpha_B = 0$), and a lowered fit relative to TNF-$\alpha$ levels, proving our equation still followed accurate biological behaviors (Fig. 2C).

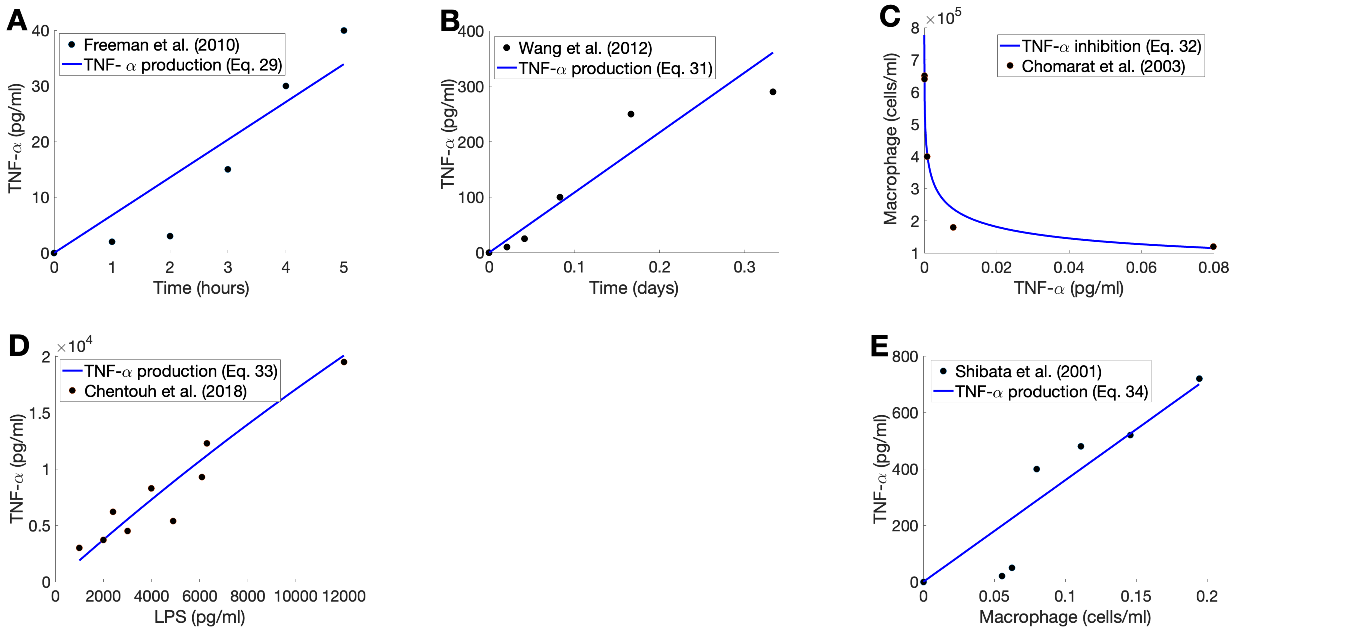


Figure 2: **TNF-$\alpha$ production by immune cells and its inhibition on the monocyte differentiation to inflammatory macrophages. A)–B)** Fitting the equation of TNF-$\alpha$ production by CD8$^+$ T cells (Eq. (29)) and NK cells (Eq. (31)), where TNF-$\alpha$ cytokine levels (pg/ml) are expressed over time (hours and days, respectively). **C)** Fitting the equation of monocyte differentiation to macrophage, induced by IL-6 while being inhibited by TNF-$\alpha$ (Eq. (32)), where macrophage production (cells/ml) is expressed over TNF-$\alpha$ cytokine levels (pg/ml) with IL-6 level fixed at 200 pg/ml. **D)** Fitting the equation of TNF-$\alpha$ production by monocytes (Eq. (33)), where TNF-$\alpha$ concentrations (pg/ml) are expressed over LPS counts (pg/ml). Lastly, **E)** Fitting the equation of TNF-$\alpha$ production by macrophages (Eq. (34)), where cytokine levels (pg/ml) are expressed over macrophage population (cells/ml).

#### *3.8. TNF-$\alpha$ production by monocytes*

When pathogens invade a host, the immune defense includes cytokine secretion by differing cells [54]. During the innate response, monocytes secrete relevant levels of TNF-$\alpha$ [54]. To discover the rate in which monocytes secrete TNF-$\alpha$ upon invasion, we fit data found in multiple sources [54, 55] to the following equation:

$$\frac{d\alpha_B}{dt} = \frac{p_{\alpha,M} M}{M + \eta_{\alpha,M}} \tag{33}$$

Fitting TNF-$\alpha$ secretion by a constant amount of monocytes over time leads us to obtain a max rate $p_{\alpha,M}$ of 110 pg/ml and a half effect $\eta_{\alpha,M}$ of 0.3851 $\times 10^9$ cells/ml (Fig. 2D).

#### *3.9. TNF-$\alpha$ production by inflammatory macrophages*

To model TNF-$\alpha$ production of inflammatory macrophages, we used the Hill function below:

$$\frac{d\alpha_B}{dt} = \frac{p_{\alpha,M_{\Phi,I}} M_{\Phi,I}}{M_{\Phi,I} + \eta_{\alpha,M_{\Phi,I}}} \tag{34}$$

The half effect is obtained by fitting the term above to temporal production data from [58], giving us $\eta_{\alpha,M_{\Phi I}} = 2.22 \times 10^{-4}$ (in $10^9$ cells/ml): see Fig. 2E. We then fit this half effect to a study by [45] which measured production of TNF-$\alpha$ under LPS stimulation. We incorporated LPS as a scalar, shown below:

$$\frac{d\alpha_B}{dt} = LPS \, \frac{p_{\alpha,M_{\Phi I}} M_{\Phi I}}{M_{\Phi I} + \eta_{\alpha,M_{\Phi I}}}. \tag{35}$$

This generates a max production rate of TNF-$\alpha$ by macrophages, $p_{\alpha,M_{\Phi I}} = 3{,}824$ pg/ml/day.

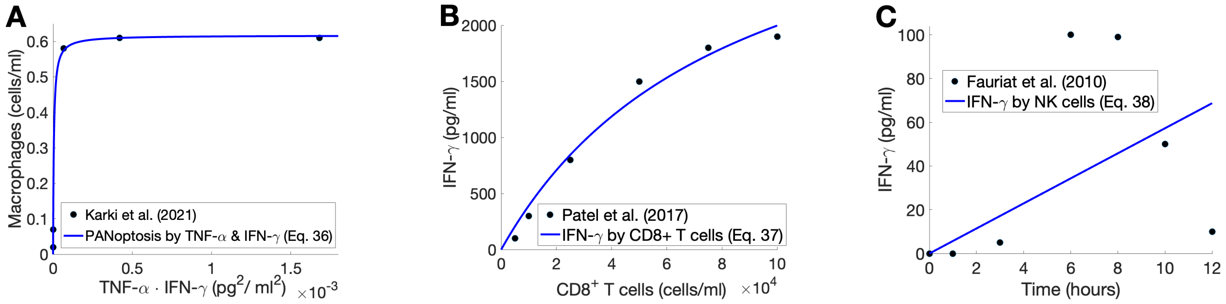


Figure 3: **Interactions between IFN-$\gamma$ and immune system.** **A)** Eq. (36) fitted to the death rate of inflammatory macrophages by varying combinations of TNF-$\alpha$ and IFN-$\gamma$: control, $2 + 1, 20 + 10, 50 + 25, 100 + 50$ ng/mL of TNF-$\alpha$ and IFN-$\gamma$, respectively [31]. This fit results in the production rate and half effect of Eq. (36). **B)** Eq. (37) was fitted to peptide stimulated production of IFN-$\gamma$. This fit was used to provide the half-effect constant of IFN-$\gamma$ production by CD8$^+$ T cells. **C)** Eq. (38) was fitted to data for IFN-$\gamma$, production by stimulated NK cells over 0, 1, 3, and 6 hours [52]. This enabled us to extrapolate the half-effect of IFN-$\gamma$ production by NK cells.

#### *3.10. Inflammatory macrophage death induced by TNF-$\alpha$ and IFN-$\gamma$*

TNF-$\alpha$ and IFN-$\gamma$ have been shown to synergistically induce programmed cell death in inflammatory macrophages [31]. We captured this behavior through the following equation:

$$\frac{dM_{\Phi I}}{dt} = -\frac{\delta_{M_{\Phi I}} \alpha_B \gamma_B}{\alpha_B \gamma_B + \epsilon_{\gamma,\alpha}}. \tag{36}$$

Fitting the equation to data found in [31] yields $\delta_{M_{\Phi I}} = 0.1543$ 1/day, and $\epsilon_{\gamma,\alpha} = 1.994 \times 10^{-5}$ (pg/ml)$^2$ (see Fig. 3A).

#### *3.11. Half-effect of IFN-$\gamma$ production by CD8$^+$ T cells*

The half-effect concentration $\eta_{\gamma,T} = 8.37 \times 10^{-5}$ (in $10^9$ cells/ml) for production of IFN-$\gamma$ was found using data from [41]. This data is fit to the following equation and the fitted curve is provided Fig. 3B.

$$\frac{d\gamma_U}{dt} = \frac{p_{\gamma,T} T}{T + \eta_{\gamma,T}}. \tag{37}$$

#### *3.12. Half-effect of IFN-$\gamma$ production by NK cells*

The half-effect concentration $\eta_{\gamma,K} = 2.99 \times 10^{-4}$ (in $10^9$ cells/ml) for production of IFN-$\gamma$ by NK cells was calculated based on data from [52]. The following equation was fitted to this data, and the fitted curve can be found in Fig. 3C.

$$\frac{d\gamma_U}{dt} = \frac{p_{\gamma,K} K}{K + \eta_{\gamma,K}}. \tag{38}$$

### 4. Parameters Calculated From Homeostasis

Remaining parameters in the model are estimated to ensure the model maintains homeostasis in the absence of infection. To do so, we require the system to return to equilibrium states after small perturbations in initial conditions for the immune cells and cytokines. Homeostasis equations are defined below, Eqs. (39)–(57), along with the corresponding parameter by setting $d/dt = 0$ in Eqs. (1)–(23). At homeostasis, we assume the absence of resistant cells and virus ($V = R = 0$). Here, $X^*$ represents homeostatic values:

$$M_{\Phi I}(0) = M^*_{\Phi I} = \frac{\left( \frac{p_{M_{\Phi I},G} {G_B^*}^{h_{M,M_\Phi}} M^*}{{G_B^*}^{h_{M,M_\Phi}} + \epsilon_{G,M_\Phi}^{h_{M,M_\Phi}}} + \frac{p_{M_{\Phi I},L} L_B^* M^*}{L_B^* + \epsilon_{L,M_\Phi}} \right)}{\left(1 - \frac{M^*_{\Phi R}}{M^*_{\Phi max}}\right) \frac{\lambda_\Phi}{\epsilon_{V,M_\Phi}} + d_{M_{\Phi I}}}, \tag{39}$$

$$F_B(0) = F_B^* = \frac{k_{B_F} T^* A_F F_U^*}{k_{int_F} + k_{B_F} F_U^* + k_{U_F}}, \tag{40}$$

$$C_B(0) = C_B^* = \frac{k_{B_C} C_U^{*\mathrm{W}_C} A_C N^*}{k_{int_C} + k_{B_C} C_U^{*\mathrm{W}_C} + k_{U_C}}, \tag{41}$$

$$C_{BF}(0) = C_{BF}^* = \frac{C_B^*}{A_C N^*}, \tag{42}$$

$$L_B(0) = L_B^* = \frac{k_{B_L}\left(T^* + N^* + M^*\right) A_L L_U^*}{k_{int_L} + k_{B_L} L_U^* + k_{U_L}}, \tag{43}$$

$$G_B(0) = G_B^* = \frac{k_{B_G} M^* A_G G_U^*}{k_{int_G} + k_{B_G} G_U^* + k_{U_G}}, \tag{44}$$

$$\gamma_B(0) = \frac{k_{B_\gamma} A_\gamma \gamma_{U,0} (M_{\Phi I,0} + T_0)}{k_{B_\gamma} \gamma_{U,0} + k_{int_\gamma} + k_{U_\gamma}} \tag{45}$$

$$\alpha_B(0) = \alpha_B^* = k_{B_\alpha}(T_{prod}^* + K(0) + M_{\mathrm{prod}}^*) A_\alpha \frac{\alpha_U^*}{(k_{int_\alpha} + k_{B_\alpha}\alpha_U^* + k_{U_\alpha})}, \tag{46}$$

$$\eta_{C,M} = \frac{p_{C,M} M^* - M^*(k_{lin_C} C_U^* + k_{B_C}(N^* A_C - C_B^*) C_U^{*W_C} - k_{U_C} C_B^*)}{k_{lin_C} C_U^* + k_{B_C}(N^* A_C - C_B^*) C_U^{*W_C} - k_{U_C} C_B^*}, \tag{47}$$

$$p_{L,M_\Phi} = \frac{M_{\Phi I}^* + \eta_{L,M_\Phi}}{M_{\Phi I}^*}\left(-\frac{p_{L,M} M^*}{M^* + \eta_{L,M}} + k_{lin_L} L_U^* + k_{B_L}\left((N^* + T^* + M^*)A_L - L_B^*\right) L_U^* - k_{U_L} L_B^*\right), \tag{48}$$

$$p_{G,M_{\Phi I}} = \frac{k_{lin_G} G_U^* + k_{B_G}(M^* A_G - G_B^*) G_U^* - k_{U_G} G_B^*}{\frac{M_{\Phi I}^*}{M_{\Phi I}^* + \eta_{G,M_\Phi}} + \frac{M^*}{M^* + \eta_{G,M}}}, \tag{49}$$

$$p_{M,G} = \frac{G_B^{*h_M} + \epsilon_{G,M}^{h_M}}{G_B^{*h_M}}\left(\frac{p_{M_\Phi,G} G_B^{*h_{M,M_\Phi}} M^*}{G_B^{*h_{M,M_\Phi}} + \epsilon_{G,M_\Phi}} + \frac{p_{M_{\Phi I},L} L_B^* M^*}{L_B^* + \epsilon_{L,M_\Phi}} + d_M M^*\right), \tag{50}$$

$$\eta_{F,M_\Phi} = \frac{p_{F,M_\Phi} M_{\Phi I}^* + \left(\frac{p_{F,M} M^*}{M^* + \eta_{F,M}} - k_{lin_F} F_U^* - k_{B_F}(T^* A_F - F_B^*) F_U^* + k_{U_F} F_B^*\right) M_{\Phi I}^*}{-\frac{p_{F,M} M^*}{M^* + \eta_{F,M}} + k_{lin_F} F_U^* + k_{B_F}(T^* A_F - F_B^*) F_U^* - k_{U_F} F_B^*}, \tag{51}$$

$$T_{prod}^* = d_T T^* - \frac{p_{T,L} L_B^* T^*}{L_B^* + \epsilon_{L,T}} - \frac{p_{T,F} F_B^* T^*}{F_B^* + \epsilon_{F,T}}, \tag{52}$$

$$p_{N,L} = N_{prod}^* = \left(d_N N^* - \frac{p_{N,L} L_B^*}{L_B^* + \epsilon_{L,N}}\right)\frac{1}{NR}, \tag{53}$$

$$p_{N,L} = N_{prod}^* = \frac{d_N N^*}{N_R + \frac{L_B^*}{L_B^* + \epsilon_{L,N}}} \tag{54}$$

$$M_{\mathrm{prod}}^* = \frac{\frac{1}{MR}\left(\frac{p_{M_{\Phi I},G} G_B^{*h_{M,M_\Phi}} M^*}{G_B^{*h_{M,M_\Phi}} + \epsilon_{G,M_\Phi}^{h_{M,M_\Phi}}} + \frac{p_{M_{\Phi I},L} L_B^* M^*}{L_B^* + \epsilon_{L,M}} + d_M M^*\right) - \psi_M^{max} \frac{G_B^{*h_M}}{G_B^{*h_M} + \epsilon_{G,M}^{h_M}}}{1 - \frac{G_B^{*h_M}}{G_B^{*h_M} + \epsilon_{G,M}^{h_M}}}, \tag{55}$$

$$K_{prod}^* = \left(p_K K_0 - \frac{p_{K,\alpha}\alpha_{B,0} K_0}{\alpha_{B,0} + \epsilon_{\alpha,K}}\right)\frac{1}{K_R}, \tag{56}$$

$$\eta_{L,M_\Phi} = \frac{M_{\Phi_I}^* - \frac{1}{p_{L,M_\Phi}}\left(-\frac{p_{L,M} M^*}{M^* + \eta_{L,M}} + k_{lin_L} L_U^* + k_{B_L}((N^* + T^* + M^*)A_L - L_B^*) L_U^* - k_{U_L} L_B^*\right) M_{\Phi_I}^*}{\frac{1}{p_{L,M_\Phi}}\left(-\frac{p_{L,M} M^*}{M^* + \eta_{L,M}} + k_{lin_L} L_U^* + k_{B_L}((N^* + T^* + M^*)A_L - L_B^*) L_U^* - k_{U_L} L_B^*\right)} \tag{57}$$

### 5. Additional Model results

#### 5.1. *All other variables for mild and severe COVID-19 disease*

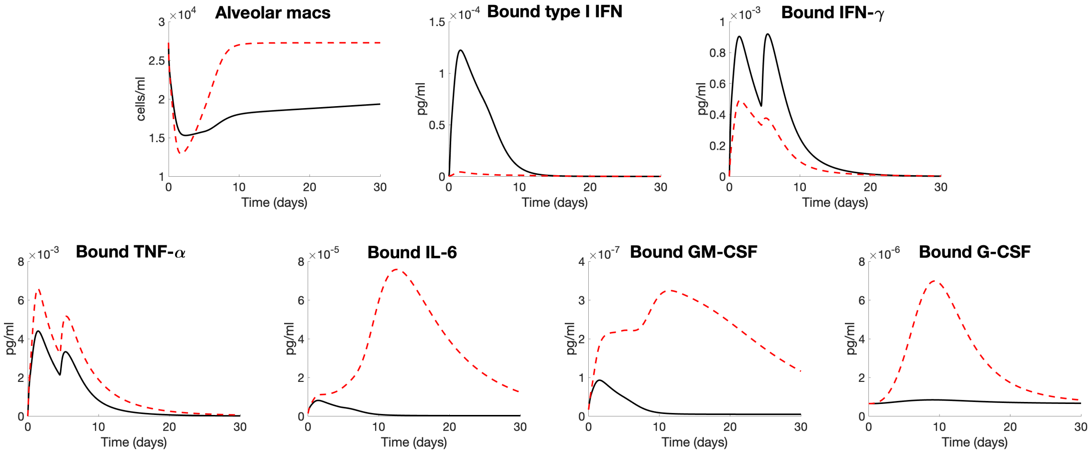


Figure 4: **Model predictions for mild vs. severe COVID-19 dynamics (for all other variables).** Mild disease (solid lines) dynamics obtained by solving Eqs. (1)–(23) with baseline parameters summarized in Tables 3–13. The graphs depict the predicted concentrations of resident macrophages and all bound cytokines over the course of a 30-day infection, and those for all other model variables can be found in Fig. 2 in the main paper.

#### 5.2. *Complete sensitivity analysis result*

See the attached heatmap from complete LSA results.

[59] R. Mehra, A. Storfer-Isser, H. L. Kirchner, N. Johnson, N. Jenny, R. P. Tracy, S. Redline, Soluble interleukin 6 receptor: a novel marker of

moderate to severe sleep-related breathing disorder, Archives of internal medicine 166 (16) (2006) 1725–1731.
[60] J. Stagg, J. H. Wu, N. Bouganim, J. Galipeau, Granulocyte-macrophage colony-stimulating factor and interleukin-2 fusion cDNA for cancer gene immunotherapy, Cancer research 64 (24) (2004) 8795–8799.
[61] P. Arnaud, Different interferons: Pharmacology, pharmacokinetics, proposed mechanisms, safety and side effects, REVUE DE MEDECINE INTERNE 23 (2002) 449S–458S.
[62] Y. Ma, S. Zhao, S. Shen, S. Fang, Z. Ye, Z. Shi, A. Hong, A novel recombinant slow-release TNF $\alpha$-derived peptide effectively inhibits tumor growth and angiogensis, Scientific reports 5 (1) (2015) 13595.
[63] J. E. Nesbitt, G. M. Fuller, Dynamics of interleukin-6 internalization and degradation in rat hepatocytes., Journal of Biological Chemistry 267 (9) (1992) 5739–5742.
[64] N. A. Nicola, L. Peterson, D. J. Hilton, D. Metcalf, Cellular processing of murine colony-stimulating factor (Multi-CSF, GM-CSF, G-CSF) receptors by normal hemopoietic cells and cell lines, Growth Factors 1 (1) (1988) 41–49.
[65] D. E. Mager, W. J. Jusko, Receptor-mediated pharmacokinetic/pharmacodynamic model of interferon-$\beta$ 1a in humans, Pharmaceutical research 19 (2002) 1537–1543.
[66] R. Fischer, R. E. Kontermann, K. Pfizenmaier, Selective targeting of TNF receptors as a novel therapeutic approach, Frontiers in cell and developmental biology 8 (2020) 401.
[67] M. Grell, H. Wajant, G. Zimmermann, P. Scheurich, The type 1 receptor (CD120a) is the high-affinity receptor for soluble tumor necrosis factor, Proceedings of the National Academy of Sciences 95 (2) (1998) 570–575.
[68] S. Tenhumberg, G. H. Waetzig, A. Chalaris, B. Rabe, D. Seegert, J. Scheller, S. Rose-John, J. Grötzinger, Structure-guided optimization of the interleukin-6 trans-signaling antagonist sgp130, Journal of Biological Chemistry 283 (40) (2008) 27200–27207.
[69] R. Sadir, E. Forest, H. Lortat-Jacob, The heparan sulfate binding sequence of interferon-$\gamma$ increased the on rate of the interferon-$\gamma$-interferon-$\gamma$ receptor complex formation, Journal of Biological Chemistry 273 (18) (1998) 10919–10925.
[70] M. Fallahi-Sichani, M. A. Schaller, D. E. Kirschner, S. L. Kunkel, J. J. Linderman, Identification of key processes that control tumor necrosis factor availability in a tuberculosis granuloma, PLoS computational biology 6 (5) (2010) e1000778.
[71] L. Snyers, V. Fontaine, J. Content, Modulation of interleukin-6 receptors in human cells, Annals of the New York Academy of Sciences 557 (1) (1989) 388–395.
[72] T. Taga, Y. Kawanishi, R. R. Hardy, T. Hirano, T. Kishimoto, Receptors for B cell stimulatory factor 2. quantitation, specificity, distribution, and regulation of their expression, The Journal of experimental medicine 166 (4) (1987) 967–981.
[73] M. Yamaguchi, M. Michishita, K. Hirayoshi, K. Yasukawa, M. Okuma, K. Nagata, Down-regulation of interleukin 6 receptors of mouse myelomonocytic leukemic cells by leukemia inhibitory factor, Journal of Biological Chemistry 267 (31) (1992) 22035–22042.
[74] S. Chiba, A. Tojo, T. Kitamura, A. Urabe, K. Miyazono, F. Takaku, Characterization and molecular features of the cell surface receptor for human granulocyte-macrophage colony-stimulating factor, Leukemia 4 (1) (1990) 29–36.
[75] D. R. Barreda, P. C. Hanington, M. Belosevic, Regulation of myeloid development and function by colony stimulating factors, Developmental & Comparative Immunology 28 (5) (2004) 509–554.
[76] A. A. Branca, Interferon receptors, In vitro cellular & developmental biology 24 (3) (1988) 155–165.
[77] S. N. Constantinescu, E. Croze, C. Wang, A. Murti, L. Basu, J. E. Mullersman, L. M. Pfeffer, Role of interferon alpha/beta receptor chain 1 in the structure and transmembrane signaling of the interferon alpha/beta receptor complex, Proceedings of the National Academy of Sciences 91 (20) (1994) 9602–9606.
[78] K. Imamura, D. Spriggs, D. Kufe, Expression of tumor necrosis factor receptors on human monocytes and internalization of receptor bound ligand, Journal of immunology (Baltimore, Md.: 1950) 139 (9) (1987) 2989–2992.
[79] E. Pocsik, R. Mihalik, F. Ali-Osman, B. B. Aggarwal, Cell density-dependent regulation of cell surface expression of two types of human tumor necrosis factor receptors and its effect on cellular response, Journal of cellular biochemistry 54 (4) (1994) 453–464.
[80] S. Kang, T. Tanaka, T. Kishimoto, Therapeutic uses of anti-interleukin-6 receptor antibody, International immunology 27 (1) (2015) 21–29.
[81] T. Hirano, IL-6 in inflammation, autoimmunity and cancer, International immunology 33 (3) (2021) 127–148.
[82] P. J. Murray, The JAK-STAT signaling pathway: input and output integration, The Journal of Immunology 178 (5) (2007) 2623–2629.
[83] R. Kurzrock, E. H. Estey, M. Talpaz, J. U. Gutterman, Granulocyte-macrophage colony-stimulating factor in myelodysplastic syndromes and aplastic anemia, Biological Response Modifiers in the Treatment of Hematopoietic Neoplasia 15 (1992) 173.
[84] MyBioSource, Recombinant Human Interferon Beta-1a (Cat. No. MBS553062), `https://www.mybiosource.com/protein/Z01001-TNF_Human.html` (2020).
[85] GenScript, TNF-$\alpha$, Human (Cat. No. Z01001), `https://www.genscript.com/protein/Z01001-TNF_Human.html` (2025).
[86] M. Rosenblum, N. Donato, Tumor necrosis factor alpha: a multifaceted peptide hormone, Critical reviews in immunology 9 (1) (1989) 21–44.
[87] J. Armstrong, E. Gluck, R. Crapo, H. A. Jones, J. Hughes, Lung tissue volume estimated by simultaneous radiographic and helium dilution methods, Thorax 37 (9) (1982) 676–679.
[88] N. Chaffey, Alberts, B., Johnson, A., Lewis, J., Raff, M., Roberts, K. and Walter, P. Molecular biology of the cell. 4th edn., Annals of Botany 91 (3) (2003) 401–401.
[89] W. Jiang, L. Kang, H.-Z. Lu, X. Pan, Q. Lin, Q. Pan, Y. Xue, X. Weng, Y.-W. Tang, Normal values for CD4 and CD8 lymphocyte subsets in healthy Chinese adults from Shanghai, Clinical and Vaccine Immunology 11 (4) (2004) 811–813.
[90] C. L. Gordon, M. Miron, J. J. Thome, N. Matsuoka, J. Weiner, M. A. Rak, S. Igarashi, T. Granot, H. Lerner, F. Goodrum, et al., Tissue reservoirs of antiviral T cell immunity in persistent human CMV infection, Journal of Experimental Medicine 214 (3) (2017) 651–667.
[91] F. Bozzano, C. Perrone, L. Moretta, A. De Maria, NK cell precursors in human bone marrow in health and inflammation, Frontiers in immunology 10 (2019) 2045.
[92] M. Osman, R. M. Faridi, W. Sligl, M.-T. Shabani-Rad, P. Dharmani-Khan, A. Parker, A. Kalra, M. B. Tripathi, J. Storek, J. W. Cohen Tervaert, et al., Impaired natural killer cell counts and cytolytic activity in patients with severe COVID-19, Blood advances 4 (20) (2020) 5035–5039.
[93] M. Maes, A.-h. Lin, L. Delmeire, A. Van Gastel, G. Kenis, R. De Jongh, E. Bosmans, Elevated serum interleukin-6 (IL-6) and IL-6 receptor

concentrations in posttraumatic stress disorder following accidental man-made traumatic events, Biological psychiatry 45 (7) (1999) 833–839.

[100] Interferon-$\gamma$ protein, recombinant humandoi:www.emdmillipore.com/US/en/product/Interferon-Protein-Recombinant-human,MM_NF-IF002.

[101] Useful pharmacokinetic equations (2013). doi:https://pharmacy.ufl.edu/files/2013/01/5127-28-equations.pdf.

[102] Interferon gamma-1bdoi:https://drugs.ncats.io/substance/21K6M2I7AG.

[103] K. Pallmer, A. Oxenius, Recognition and regulation of T cells by NK cells, Frontiers in immunology 7 (2016) 251.

[104] L. Cifaldi, G. Prencipe, I. Caiello, C. Bracaglia, F. Locatelli, F. De Benedetti, R. Strippoli, Inhibition of natural killer cell cytotoxicity by interleukin-6: implications for the pathogenesis of macrophage activation syndrome, Arthritis & rheumatology 67 (11) (2015) 3037–3046.

[105] P. Chomarat, J. Banchereau, J. Davoust, A. Karolina Palucka, IL-6 switches the differentiation of monocytes from dendritic cells to macrophages, Nature immunology 1 (6) (2000) 510–514.
