## Supplementary figures and images for "Immune Cell Dysfunction of SARS-CoV-2: Mathematical Modeling of the Within-Host Immune Dynamics"

### Complete Sensitivity Analysis Results

#### 5% increase in parameter value

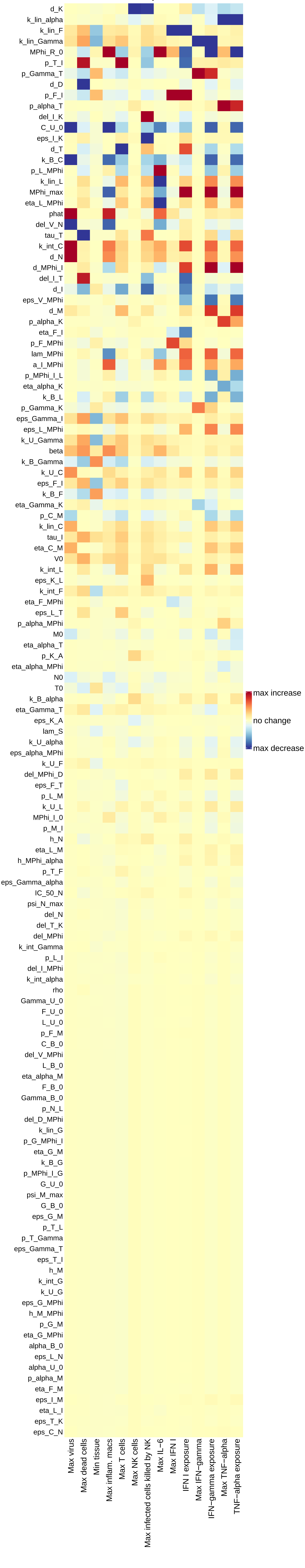
